# Partially Shared Multi-Modal Embedding Learns Holistic Representation of Cell State

**DOI:** 10.1101/2024.10.01.615977

**Authors:** Xinyi Zhang, GV Shivashankar, Caroline Uhler

## Abstract

Experimental technologies for jointly measuring different data modalities at the single-cell level offer different windows into cell state. To obtain a holistic understanding of cell state, computational methods are needed that carefully integrate the different views to capture shared information as well as tease apart modality-specific information. We present a computational framework that automatically learns partial information sharing between multiple modalities by using an **A**utoencoder with a **P**artially **O**verlapping **L**atent space learned through **L**atent **O**ptimization (APOLLO). On paired scRNA-seq and scATAC-seq data (SHARE-seq) and paired scRNA-seq and surface protein data (CITE-seq), we demonstrate that APOLLO comprehensively and automatically identifies and distinguishes between information captured by both modalities, in the shared latent space, and modality-specific information. Beyond sequencing modalities, large-scale multiplexed single-cell imaging datasets, such as the Human Protein Atlas, are becoming available that allow studying how protein localization relates to function. While chromatin, microtubules or ER are standardly stained as a reference, little is known about the information shared between these stains. We found that APOLLO enables the prediction of missing modalities, such as unmeasured protein stains, and allows disentangling which modality or cellular compartment is linked with a specific phenotype, such as the variability in protein localization observed across single cells. Collectively, APOLLO enables explicit learning of shared and modality-specific information leading to a more holistic understanding of cell state and the underlying regulatory mechanisms. APOLLO is a general framework that can be applied to any multi-modal data well beyond the single-cell domain including, for example, large-scale medical biobanks.

## Introduction

Single-cell technologies allow studying the cell state changes and regulatory mechanisms underlying development, disease, and other biological processes. While each data modality offers a different view, the true underlying cell state remains not directly observable, and different technologies have been developed to reveal a particular aspect of a cell’s state. For example, while chromatin imaging captures the physical state of a cell, transcriptomic measurements provide an integrated view on nuclear signaling outputs, and multiplexed imaging or epitope sequencing of proteins reflects the functional state of a cell. Importantly, given that the physical state of a cell, nuclear signaling, and the functional state of a cell are correlated, we expect some information to be shared among the different modalities, while some aspects of cell state are only captured in one of the modalities (Figure 1a). The development of experimental and computational technologies for jointly measuring and carefully integrating different data modalities to identify shared as well as modality-specific information is thus critical to obtain a holistic understanding of cell state and the underlying regulatory mechanisms.

**Figure 1.**
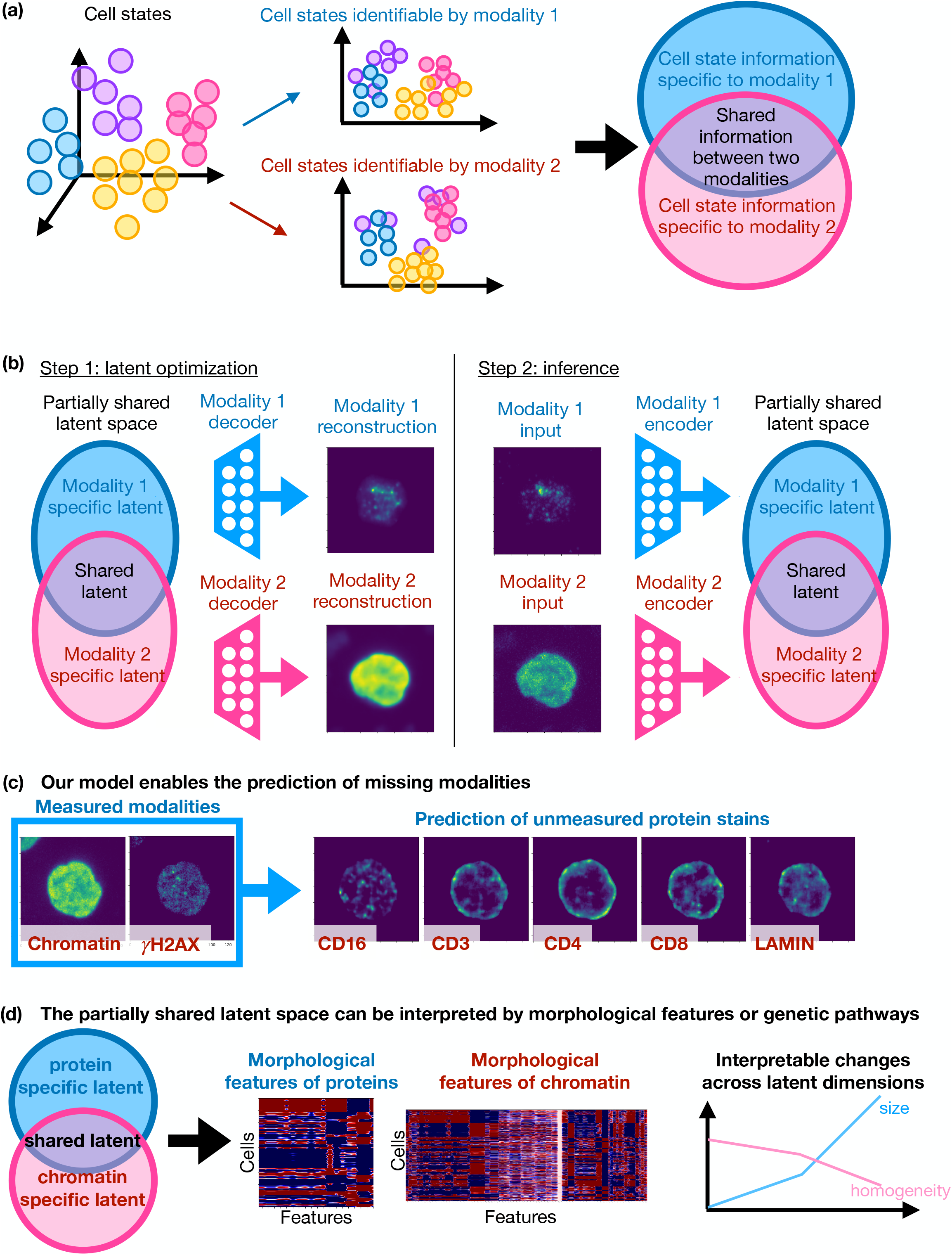
APOLLO enables learning of partial information sharing across data modalities and provides a framework for cross-modality prediction. (a) Since different experimental technologies capture different aspects of cell state, we expect some cell state information to be shared among different modalities and some information to be modality-specific. (b) APOLLO is a novel autoencoder-based approach that learns three latent spaces to disentangle information captured by each modality. APOLLO uses a two-step training procedure. In step 1, the decoders of each modality are trained so that the decoders can reconstruct the input data from the latent spaces. In step 2, modality-specific encoders are trained to enable inferring the latent space embedding for cells not used in training the model. (c) The explicit learning of partial information sharing allows APOLLO to perform accurate cross-modality prediction. The example shows the predicted protein (CD16, CD3, CD4, CD8, lamin) fluorescent images of a cell in a held-out patient, given the cell’s chromatin and *γ*H2AX protein stain. (d) The information captured by the shared and modality-specific latent spaces learned by APOLLO is interpretable.

Recent advances in experimental techniques have enabled the simultaneous measurement of multiple modalities, including multiplexed imaging and different sequencing modalities at single-cell level [1, 2, 3, 4, 5, 6], and various spatial transcriptomic methods in the tissue context [7, 8, 9]. Current computational methods for analyzing multi-modal data mostly process each modality separately and then compare the results in a second step. For example, recent work in the context of large-scale multiplexed imaging separately extracted image features of each protein stain and performed separate hypothesis tests in pre-defined manually annotated classes to infer metabolic traits [10]. Such analysis is restricted to features identified in prior work and cannot exploit the information in the multi-modal dataset to identify morphological changes across cellular components underlying a cell state change. Analyzing each modality separately in multi-modal data also means that comparison and interpretation of information contained in the different modalities has to be made manually, which usually restricts the analysis to a few cell clusters or features [5, 10]. In the context of paired scATAC-seq and scRNA-seq data, chromatin accessibility is often summarized at the gene-level in order to allow direct comparison to gene expression [5]. Such coarse-graining of the data into a shared set of features may lose important information in each modality and can only be done when the modalities are similar. To overcome this limitation, representation learning methods [11] have been introduced to single-cell biology [12, 13, 14] enabling the automatic integration of multi-modal data for joint downstream analysis including clustering and biomarker identification [15, 16, 17, 18, 19, 20, 21]. For example, in prior work we developed approaches based on autoencoders, popular neural networks used for representation learning, to extract features from each modality and embed the features of all input modalities into a shared latent space for downstream analysis; we applied this framework to chromatin imaging, RNA-seq and ATAC-seq in the single-cell domain, as well as spatial transcriptomics [15, 16]. However, existing multi-modal integration methods learn the union of information from the input modalities, incorporating both the shared and modality-specific information, without disentanglement [16, 22, 19]. Given the explosion of large-scale paired measurements in the sequencing and imaging domain at the single-cell and tissue level but also in the medical domain in large-scale biobanks that collect paired information on many individuals [23, 24], computational methods are needed that automatically and comprehensively learn the shared and modality-specific information to fully exploit multi-modal measurements.

We present a method that automatically learns partial information sharing between multiple modalities by using an **A**utoencoder with a **P**artially **O**verlapping **L**atent space learned through **L**atent **O**ptimization (APOLLO). APOLLO is motivated by recent theoretical and algorithmic developments: (1) Under simplifying assumptions, in particular linearity of decoders and non-Gaussian distribution of each modality, recent theoretical work showed that the shared and modality-specific latent representation is provably identifiable and that the identified features are causal [25], thereby indicating that the considered problem of identifying shared as well as modality-specific information from multi-modal data is well-posed and that disentangling information in this way may lead to more interpretable/causal features. (2) Recently, a novel two-step training strategy was introduced, which first learns the latent embedding together with the decoder (i.e., map from the latent embedding to a given data modality) and then learns the encoder (i.e., map from a given data modality into the latent space) [26]. Such latent optimization strategy was shown to successfully disentangle latent spaces corresponding to image class and content in well-characterized cars, poses, and faces datasets. APOLLO builds on these two recent developments to explicitly learn a latent space that represents information that is shared between modalities and a latent space per modality containing information that is modality-specific (Figure 1b). As a proof-of-concept, we demonstrate using a paired scRNA-seq and scATAC-seq dataset (SHARE-seq [5]) that APOLLO can comprehensively and automatically identify and distinguish between gene activity captured by both, transcriptomics and chromatin accessibility, as well as by only one of the modalities. We further apply APOLLO to a paired scRNA-seq and surface protein dataset (CITE-seq [17]) to demonstrate that APOLLO can correctly disentangle known shared and modality-specific information. Beyond sequencing modalities, large-scale multiplexed single-cell imaging datasets are becoming available that allow studying how protein localization relates to function. While chromatin staining is standardly used in these datasets as a reference and chromatin staining has been shown to contain rich cell state information, little is known about the information shared between chromatin organization and protein localization, reflecting the regulation and signaling between chromatin and proteins. APOLLO identifies features of chromatin organization and protein localization that correspond to cell state information shared between the two modalities as well as morphological features that are only captured by one modality. The explicit learning of partial information sharing by APOLLO also enables accurate cross-modality predictions, such as predicting unmeasured proteins from chromatin imaging (Figure 1c). Incorporating additional cellular stains, including microtubule and ER, we demonstrate on multiplexed imaging data from the Human Protein Atlas [27] that APOLLO can be used to uncover novel associations between variations in protein subcellular localization and the morphology of different cellular compartments. Overall, we demonstrate that APOLLO provides a general framework that enables disentangling and interpreting the shared and modality-specific information contained in multi-modal datasets to obtain novel biological insights by learning partially shared latent spaces (Figure 1d).

## Results

### 1 APOLLO enables learning of partial information sharing across data modalities

To explicitly model shared and modality-specific information, we propose a new framework and an accompanying training strategy to represent partial information sharing between modalities as partially overlapping latent spaces (Figure 1b). While a regular autoencoder learns a representation of a single modality, our previous works have demonstrated that a joint representation of multiple modalities can be obtained by using one autoencoder per modality and aligning their latent spaces [15, 16]. However, such alignment of the latent spaces results in a single latent space that entangles shared and modality-specific information. To overcome this limitation, APOLLO only enforces alignment between a subset of latent dimensions and allows the remaining latent dimensions to account for domain-specific information.

More specifically, let *Z* = (*Z*_*i*_)_*i*∈*H*_ be a latent random vector with distribution *P*_*Z*_ representing the true underlying state of a cell. Each single-cell technology may only measure some aspect of cell state. Let *H*_*j*_ ⊆ *H* represent the subset of latent variables that can be accessed by technology *j* and let *D*_*j*_ be a modality-specific map, which takes in the latent vector 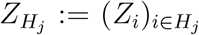 and outputs a data sample *X*^*(j)*^ from modality *j*, i.e., 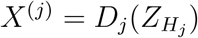 Note that *X*^(*j*)^ may be high-dimensional (e.g., 20,000-dimensional for single-cell RNA-seq technologies or consisting of the number of pixels in a chromatin image). Assuming that we have access to 1 ≤*j* ≤*J* paired modalities that can be measured on the same cell, then each cell gives rise to samples (*X*^(1)^,…, *X*^(*J*)^) from a joint distribution *P* (*X*^(1)^,…, *X*^(*J*)^). Given such samples, the problem then becomes to identify the latent features *Z* and in particular understand which latent features are shared across modalities and which are modality-specific; in the case of *J* = 2, this leads to the task of identifying the shared features *Z*_*S*_ = *Z*_*H*1⋂*H*2_ as well as the modality-specific features 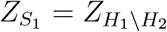 and 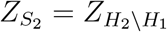 To generalize to *J >* 2, *Z*_*S*_ could represent the shared information across all modalities, i.e. 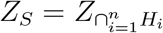 and 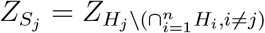

Without additional assumptions, the shared and modality-specific latent features may not be identifiable. For linear maps *D*_*j*_, a recent theoretical study proved identifiability under additional assumptions, such as non-Gaussianity of each marginal distribution [25]. Furthermore, when parameterizing the maps *D*_*j*_ with neural networks and using a latent optimization strategy for training, a recent study demonstrated empirically in the context of computer vision that simple image aspects could be disentangled and represented by different latent features [26]. APOLLO builds on these works to automatically learn partial information sharing between multiple modalities by using an autoencoder framework with a partially overlapping latent space learned through latent optimization. While our framework is general, for simplicity we describe it in the bi-modal setting where *J* = 2. In this context, *D*_1_ and *D*_2_ are decoders that map from shared and modality specific latent features to each data modality. Given data (*x*^(1)^, *x*^(2)^) from *P* (*X*^(1)^, *X*^(2)^), the corresponding shared latent features as well as the modality-specific features are learned simultaneously with the decoders *D*_1_ and *D*_2_ by minimizing the reconstruction error 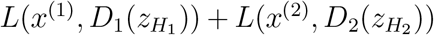 where *L*(·) is the mean squared error or another appropriate loss function. In practice, the dimension of the shared latent space (i.e. *S* := *H*_1_ ⋂ *H*_2_) is chosen much larger than the dimension of the modality-specific latent spaces (i.e., *S*_1_ := *H*_1_ \*H*_2_ and *S*_2_ := *H*_2_ \*H*_1_), to enforce that information shared by the two modalities is represented in the shared latent space. To encourage this further as well as to enable cross-modality prediction (described below), we introduce two additional decoders 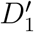 and 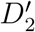 that map from the shared latent space to each of the modalities, trained by minimizing the reconstruction error 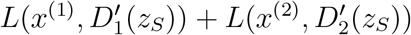 In the applications of APOLLO to the SHARE-seq and multiplexed imaging datasets described in the following sections, we demonstrate that APOLLO is robust to the choice of latent space dimensions as well as to whether the two additional decoders 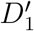 and 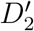 are included. In practice, similar to variational sampling in autoencoders, in order to improve generalization to unseen samples, we add Gaussian noise to each component of the latent space and add a regularization term (*l*_2_-norm of the latent features). Thus the full objective function is:

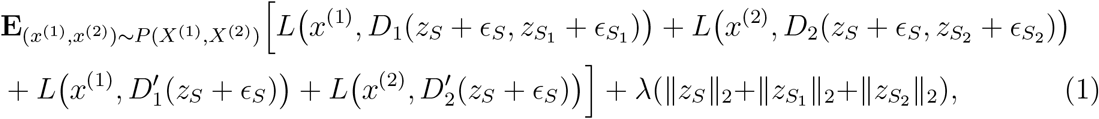

where ∈_*s*_, 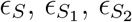 denote Gaussian noise in each component of the latent space and λ is a hyperparameter. We minimize (1) to obtain the decoders *D*_1_, *D*_2_, 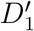 and 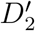 as well as the shared latent features *z*_*S*_ and the modality-specific latent features 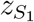 and 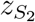

In order to generalize to unseen samples and to enable cross-modality prediction, we use a second training step to obtain encoders *E*_*j*_ that map from each data modality to their respective latent spaces. More precisely, given a sample (*x*^(1)^,…, *x*^(*J*)^) and the corresponding features *z* obtained from the first training step, the encoders *E*_*j*_, 1≤ *j* ≤ *J*, are obtained by minimizing the mean-squared error 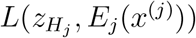 The encoders and the previously described decoders can be any neural network architecture that is suitable for the particular data modality, such as convolutional networks for images and fully connected networks for gene expression. For cross-modality prediction, a sample *x*^(*j*)^ from input modality *j* is first encoded to the latent space to obtain 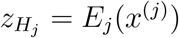 Then the decoder 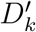 of the target modality *k* is applied to the dimensions corresponding to the shared latent space to obtain 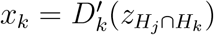 Additional details of model setup and training are provided in SI Appendix, Extended Methods.

In the following, we demonstrate that APOLLO is generally applicable to any multi-modal data by discussing four different multi-modal settings, including paired sequencing-based modalities as well as multiplexed imaging.

### 2 APOLLO provides a general framework for the integration of paired sequencing-based measurements

We demonstrate that APOLLO learns meaningful shared and modality-specific latent spaces and is generally applicable to paired sequencing-based measurements using two representative datasets: paired scRNA-seq and scATAC-seq measured using SHARE-seq [5] (SI Appendix, Figures. S1 and S2) and paired scRNA-seq and cellular surface protein abundance measured using CITE-seq [17] (SI Appendix, Figure S3a).

Using the SHARE-seq data, we first assess if the modality-specific latent spaces capture additional information compared to the shared latent space using a cell type classification task (SI Appendix, Extended Methods). As a baseline, we used the cell types defined by Ma *et al*. to train a classifier that predicts the cell types using the shared latent space. Additionally, we trained two separate classifiers that use the ATAC-seq latent space and RNA-seq latent space respectively, in addition to the shared latent space. To ensure that our model is robust to the choice of latent space dimensions, we tested different dimensions for the shared and modality-specific latent spaces as well as different ratios between the latent space dimensions (Figure 2a). The incorporation of either the RNA-specific latent space or the ATAC-specific latent space improves the cell type classification accuracy compared to using the shared latent space alone for all latent space dimensions. This shows that the modality-specific latent spaces can capture biologically meaningful information that is not represented in the shared latent space, and our model is robust to the choice of latent space dimensions.

**Figure 2.**
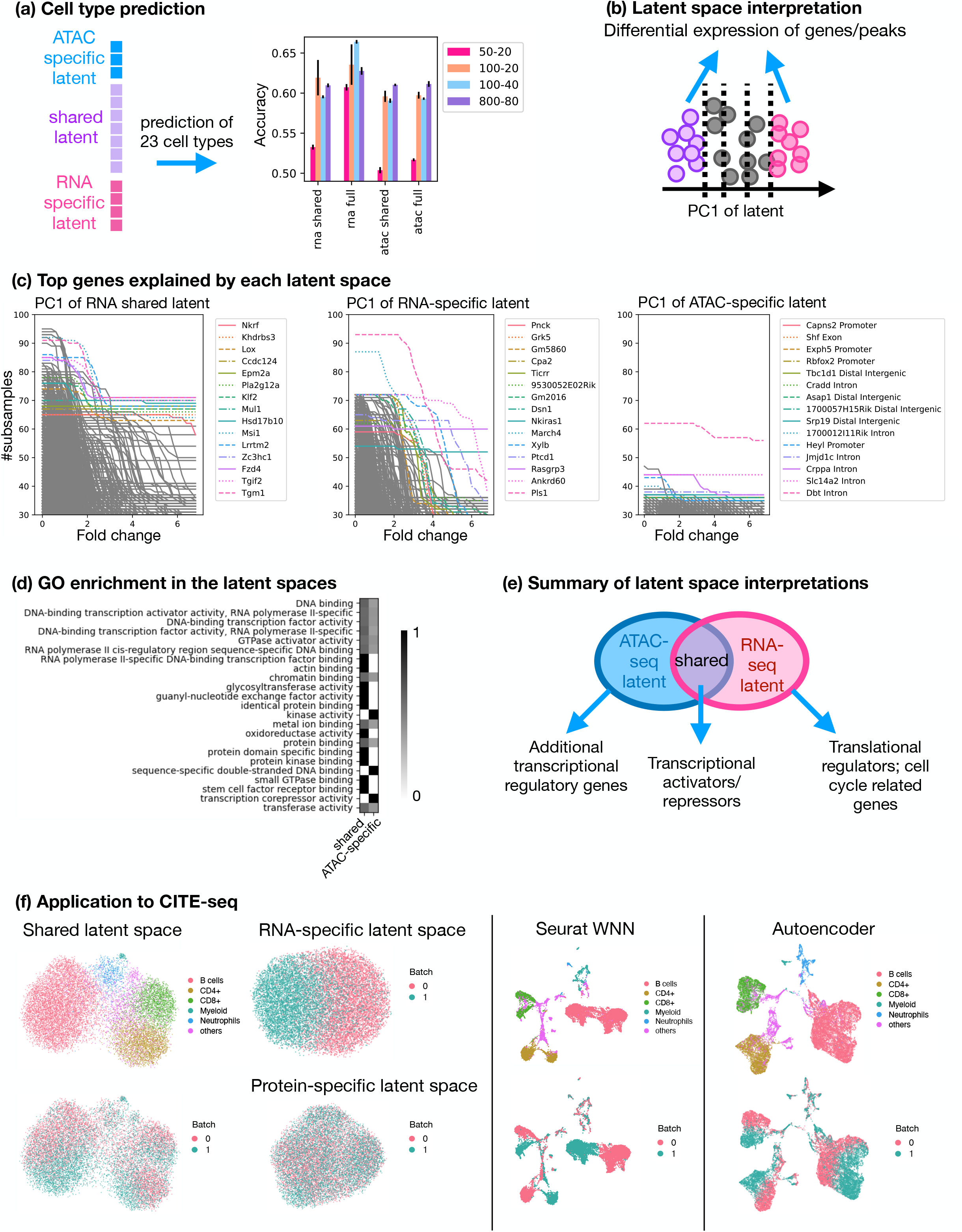
APOLLO enables the identification of shared and modality-specific information in paired scRNA-seq and scATAC-seq data. (a) Four separate models with different latent space dimensions are tested. The dimensions are listed as “shared latent space dimension”-”modality-specific dimension”. For each model, we train four separate classifiers, with three random train-test splits, that predict cell type based on the following inputs respectively: 1) the shared latent space encoded using scRNA-seq (rna shared) or scATAC-seq (atac shared), 2) the shared latent space concatenated with the RNA-specific latent space (rna full) or the atac-specific latent space (atac full). The plot shows the mean and standard deviation of each model. The baseline accuracy computed by randomly permuting cell type assignments is 0.108 with a standard deviation of 0.03 across 5 random permutations. (b)-(c) The genes or peaks explained by each latent space can be identified by performing differential expression using the cells at the two ends and at the center of each principal component (PC, panel b). The curves in (c) show the number of times among 108 random train-test splits that a gene passes a particular fold-change threshold, while the p-value cutoff is set to 0.05 after multiple testing correction (SI Appendix, Extended Methods). The top genes with the highest area under the curve are colored. (d) We performed gene ontology (GO) enrichment analysis using the DE genes of the top PCs identified as described in (c). The enriched GO molecular function terms in the shared and ATAC-specific latent spaces are shown. The heatmap shows the proportion of times a GO term is observed to be enriched in one of the latent spaces. (e) Summary of genes captured in each of the three latent spaces. (f) Application of APOLLO to the CITE-seq data from [17] shows correct disentanglement of shared and modality-specific information. Left panel: UMAPs of the shared and the two modality-specific latent spaces. Middle panel: UMAPs obtained from Seurat WNN analysis. Right panel: UMAPs obtained from the shared latent space of standard multi-modal autoencoder with an encoder and a decoder for each input modality (SI Appendix, Extended Methods). Both UMAPs show the shared latent space encoded using the protein encoder. The cells are colored by cell types or experimental batches.

We next demonstrate how principal component analysis (PCA) can be used to interpret the information contained in the shared and modality-specific latent spaces. Along each principal component (PC), we bin the cells based on their positions along the PC and then compare the gene expression or peak counts of the cells at the two ends of the PC and at the center of the PC (Figure 2b, SI Appendix, Extended Methods). We identify cell cycle related genes among the top genes explained by the first PC of the RNA-specific latent space, such as Ticrr and Dsn1 [28, 29] (Figure 2c), while the top PCs of the ATAC-specific latent space capture the activity of promoters of transcriptional regulators, e.g. the promoter of the transcription factor Heyl (Figure 2c and SI Appendix, Figure S4b). Furthermore, the top genes explained by the first three PCs of the RNA and ATAC shared latent space contain known transcriptional regulators, some of which are known transcriptional activators or repressors (e.g. *Zeb1*) based on the correlation between transcription factor (TF) activity and TF RNA expression levels [5] (Figure 2c and SI Appendix, Figure S5a). Interestingly, the top genes of the ATAC-specific or the shared latent space are generally not related to cell cycle and the top genes of the RNA-specific latent space are not related to transcriptional regulation; exceptions are the gene Nkrf captured in the shared latent space and the gene CBx2 captured in the RNA-specific latent space, which are known to regulate cell cycle [30, 31]; see Figure 2c and SI Appendix, Figure S5b.

Gene ontology (GO) enrichment analysis on the genes explained by each latent space also identifies GO terms related to transcriptional regulation in the shared latent space and the ATAC-specific latent space but not in the RNA-specific latent space (Figure 2d). This could indicate a time lag between the change in chromatin accessibility and the resulting change in gene expression. Interestingly, GO terms related to post-transcriptional regulation are found in the shared latent space. These analyses demonstrate that the shared and modality-specific latent spaces identified by APOLLO are biologically meaningful and can be efficiently analyzed to interpret the shared and modality-specific information (Figure 2e).

To further demonstrate that APOLLO can correctly disentangle shared and modality-specific information for different kinds of sequencing-based measurements, we applied APOLLO to a CITE-seq dataset of murine spleen and lymph nodes with paired scRNA-seq and surface protein measurements obtained from two wild-type mice processed in two separate experiments [17]. We used the same training strategy and model architecture as in the application to the SHARE-seq data. Since cross-modality prediction isn’t the goal in this application, we removed the two decoders 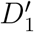 and 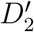 that map from the shared latent space to each of the modalities (SI Appendix, Figure S3a). Using a standard preprocessing and visualization procedure implemented in Scanpy [32], the resulting UMAP plots show that the major cell types can be separated by either scRNA-seq or protein abundance and that the scRNA-seq data shows further separation of the cells by mouse, indicative of experimental batch effects (SI Appendix, Figure S3b-c). The latent spaces of our APOLLO model correctly disentangle the multi-modal information: the shared latent space between scRNA-seq and protein abundance shows separation by cell types but not by experimental batches, while batch separation is captured by the modality-specific latent space of scRNA-seq (Figure 2f). In contrast, applying existing multi-modal integration methods without additional batch correction, such as the popular weighted-nearest neighbor (WNN) method in Seurat [22] or a standard multi-modal autoencoder [33], results in a latent space that learns the union of information from both modalities, in which cells are separated by both cell types and batches without disentanglement (Figure 2f and SI Appendix, Figure S3d and Extended Methods). This analysis further demonstrates that APOLLO correctly disentangles the shared and modality-specific information while integrating different kinds of paired-sequencing modalities, which could not be achieved by existing methods that only perform integration.

### 3 APOLLO learns partially shared latent spaces of chromatin and different proteins through conditioning

While so far we demonstrated an application of APOLLO to learn the partially shared latent spaces between paired sequencing-based modalities, APOLLO is a general framework that can be applied to any paired multi-modal data by choosing the appropriate modality-specific encoders and decoders. Furthermore, APOLLO can directly be applied to integrate more than two modalities. We introduce a version of APOLLO that incorporates conditioning in the latent space to allow for the integration of chromatin images and images of multiple protein types.

In the following, we apply APOLLO to a multiplexed imaging dataset of human peripheral blood mononuclear cells (PBMCs) from [34]. The dataset consists of a total of 32,345 PBMCs from 40 patients with one of four diagnoses: healthy, meningioma, glioma, head and neck tumor (SI Appendix, Figure S6). For each patient, two different multiplexed imaging datasets were obtained, namely one subset of cells was stained for CD4, CD8, CD16, and DAPI, while the other subset of cells was stained for Lamin, CD3, *γ*H2AX, and DAPI. To incorporate multiple proteins, a trainable vector of protein ID, shared across all images of the same protein, is concatenated to the inputs to the decoders as well as to the last hidden layer of the encoders (SI Appendix, Figure S7-8 and Extended Methods). This conditional model accurately reconstructs images of cells in held-out individuals (Figure 3a and SI Appendix, Figure S9), which ensures that APOLLO can comprehensively capture biologically relevant information from the input modalities.

**Figure 3.**
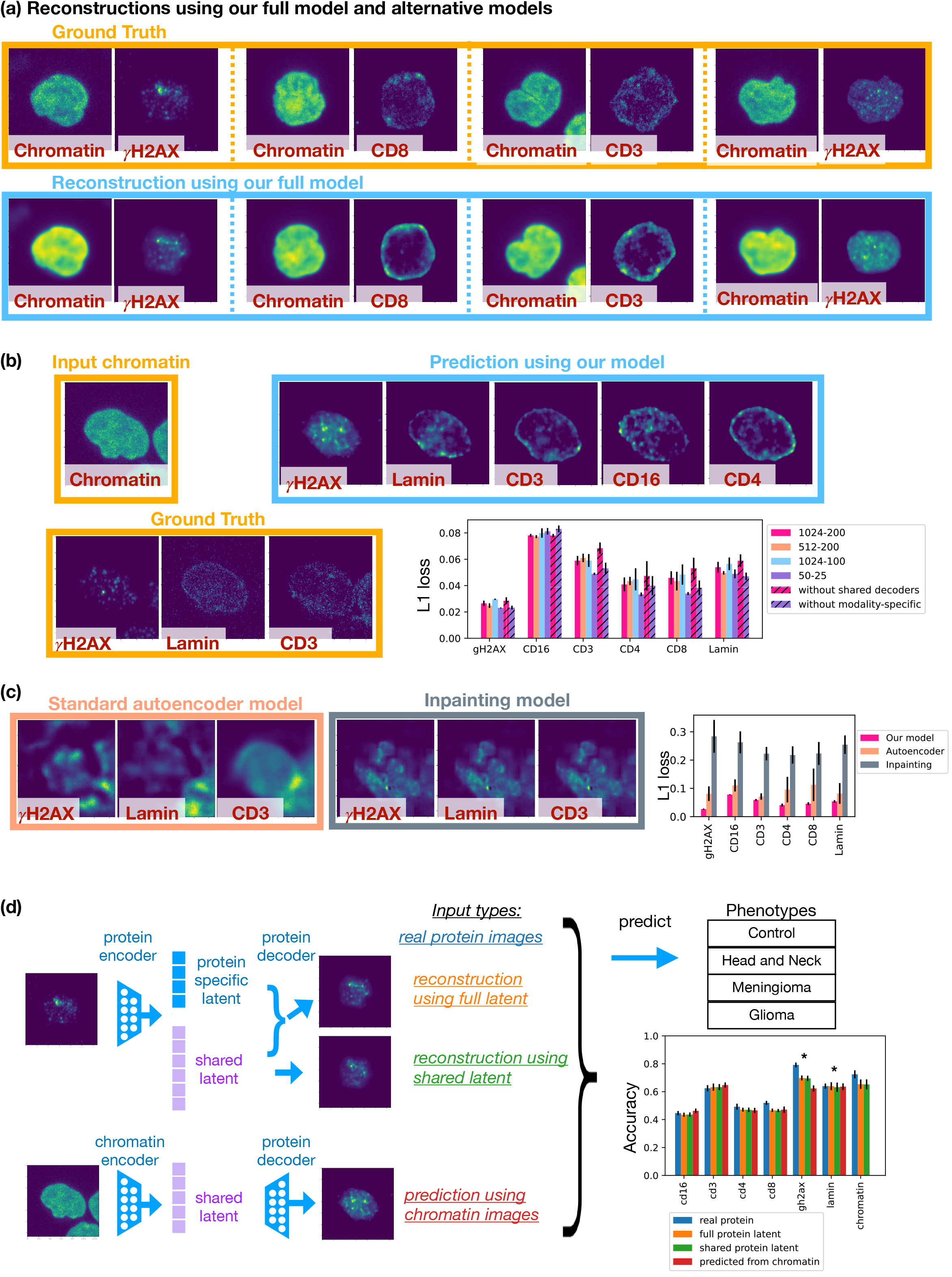
APOLLO enables accurate prediction of protein localization from chromatin imaging. (a) Examples are shown of reconstructed images of cells from held-out patients. (b) Protein predictions from chromatin images are shown for a cell where the ground truth protein images are available for the three proteins (*γ*H2AX, lamin, CD3). The cell was randomly selected among all cells with relatively good ground truth images in all three protein channels. The average prediction loss for each protein is quantified comparing the APOLLO model with different latent space dimensions to two alternative models, namely our model without the decoders that map from the shared latent space to the output (without shared decoders) and our model without separating the shared and modality-specific latent spaces (without modality-specific). The different latent space dimensions tested are listed as “shared latent space size”-”modality-specific latent space size”, e.g., 1024-200. Each model is trained with at least 4 different random initializations and the error bars show the standard deviations. (c) Cross-modality predictions obtained using our full model are compared to those obtained using a standard multi-modal autoencoder training procedure as well as a prior image inpainting model [36] (SI Appendix, Extended Methods). Each model is trained with at least 4 different random initializations and the error bars show the standard deviations. Protein predictions from chromatin images are shown for the same cell used in (a). (d) We compare the performance of using real protein/chromatin images (real protein), the reconstructed images from the full latent space (full protein latent), the reconstructed images from only the shared latent space (shared protein latent), and the protein images predicted from chromatin images (predicted from chromatin). Error bars show the result of different train-test splits. *: The prediction performance of the protein images reconstructed from the full latent space is significantly better than the protein images reconstructed from only the shared latent space (p-value = 0.0011 for *γ*H2AX and p-value = 0.098 for lamin).

### 4 APOLLO enables accurate cross-modality prediction of protein localization from chromatin imaging

The number of proteins that can be simultaneously imaged in a cell is limited, ranging from a single protein using endogenous tagging [35] to around 30 proteins in fixed samples [1]. In the following, we show that the shared latent space learned by APOLLO enables cross-modality predictions and can be applied to predict the unmeasured proteins in a cell from the chromatin image of that cell (SI Appendix, Figure S7a and Extended Methods).

The task of predicting images of unmeasured protein stains has been considered using supervised neural network models [36, 37]. In particular, image inpainting methods have been applied to predict the protein image of a target cell from its chromatin and microtubule stains given all three images of other cells [36]. While such inpainting methods can capture the average distribution, they do not produce realistic single-cell images (Figures 3b, 3c, and SI Appendix, Figures S10e, and S11). In contrast, APOLLO can accurately predict unmeasured proteins at the single-cell level in held-out individuals based on chromatin images (SI Appendix, Figure S11), and the performance of APOLLO is robust to the choice of latent space dimensions (Figure 3b-c). To interrogate the necessity of each component of APOLLO, we perform an ablation study for the cross-modality prediction task. We first compare our two-step latent optimization training to the standard autoencoder training procedure, where encoders and decoders have the same architecture as our default APOLLO model but are trained simultaneously without directly updating the parameters of the latent spaces (SI Appendix, Figures S10a and S12a). We find that standard one-step training is not able to generate realistic protein images in held-out individuals (Figures 3b-c,and SI Appendix, Figure S11). Furthermore, we tested the effect of removing the separation between the shared and modality-specific latent spaces while keeping the same two-step training procedure (SI Appendix, Figures S10b-d, S12b). This model without disentanglement of modality-specific information shows comparable performance to our APOLLO model that uses only the (smaller-dimensional) shared latent space for prediction (Figure 3b), which indicates that the shared latent space has captured all shared information. These results suggest that the improvement in prediction performance by APOLLO is mainly a result of the two-step training procedure and that the latent space disentanglement allows for the use of a smaller latent space dimension (2/3 of the full latent space) for cross-modality prediction at test time. Similar as in the applications to paired-sequencing modalities, it is also possible to remove the decoders that map from the shared latent space to each of the modalities while maintaining comparable performance in cross-modality prediction (Figure 3b).

Next, we demonstrate that the protein images predicted from chromatin images have similar performance on a downstream classification task as the real protein images. For each protein, we train four separate convolutional neural network classifiers with the same architecture to predict the phenotype of the individual from whom the input cell image was obtained from: healthy, meningioma, glioma, or head and neck tumor. For each protein, the four classifiers use the following inputs respectively: 1) real protein images; 2) reconstructed protein images using the full protein latent space, i.e., the shared latent and protein-specific latent spaces; 3) reconstructed protein images using only the shared latent space; 4) protein images predicted from chromatin images through the shared latent space. While the reconstructed protein images have similar phenotype classification accuracy across different proteins as the real protein images (Figure 3d), reconstruction from the full latent space has slightly better classification accuracy than reconstruction from the shared latent space for *γ*H2AX and lamin (Figure 3d), thereby indicating that the protein-specific latent spaces are able to learn disease-relevant information that is not shared by chromatin. Moreover, Figure 3d shows that the predicted protein images and the real protein images have comparable phenotype classification accuracy, indicating that CD3 is a better predictor of phenotype than CD16, CD4, or CD8, which suggests that the protein images predicted from chromatin capture similar disease-relevant information as the real protein images. These results demonstrate that APOLLO is capable of accurately predicting protein localization from chromatin organization and the predicted protein images can be used for downstream tasks with performance mimicking that of real protein images.

### 5 APOLLO identifies interpretable morphological features that capture the shared and modality-specific information between chromatin organization and protein localization

In contrast to paired sequencing measurements, where the different modalities can be compared and interpreted at the level of genes [5], shared features are not directly available for paired image modalities. APOLLO provides a systematic framework that can be applied also in these settings to interrogate the information that is shared between modalities and specific to each modality. To identify the morphological features captured by the shared and modality-specific latent spaces, we perform PCA and bin cells along each PC (SI Appendix, Extended Methods). Cells along a particular PC can then be sampled and visualized to interrogate the information captured in the shared or modality-specific latent spaces (Figure 4a). We use the set of chromatin and protein features defined in [34] to compare the cells at the two ends of each PC and at the center of the PC for each latent space, thereby allowing us to identify the main features captured by each latent space (Figure 4a and SI Appendix, Figures S13-19, Extended Methods). For example, we find that the first PC of the shared latent space mainly captures *bounding box area* and *homogeneity of chromatin* (Figure 4a).

**Figure 4.**
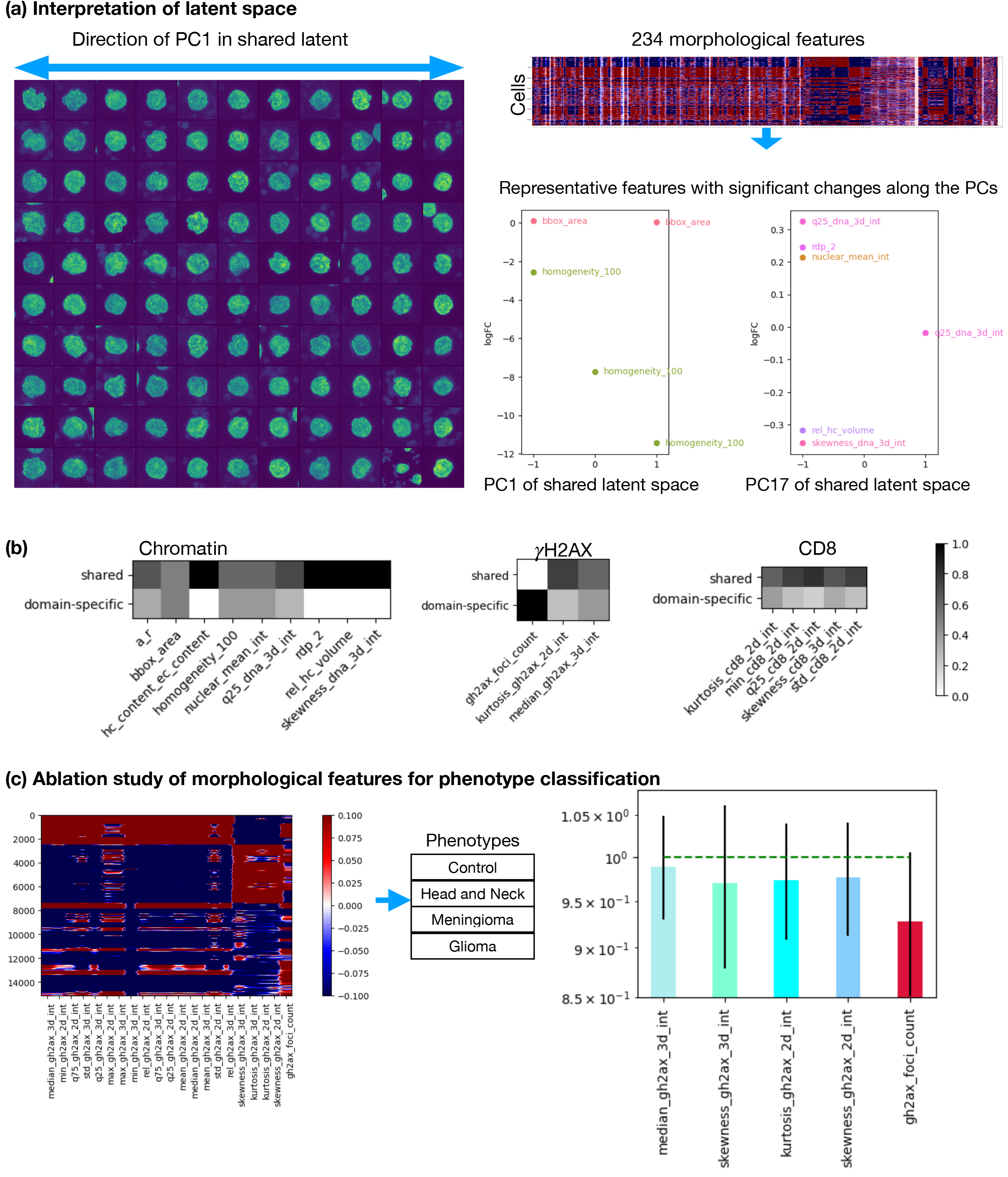
APOLLO identifies interpretable morphological features captured in the shared and modality-specific latent spaces of chromatin organization and protein localization. (a) Left: Cells are separated along the first PC of the chromatin shared latent space into 11 bins of equal percentiles; randomly selected cells in each bin are shown. Right: 234 predefined handcrafted morphological features from [34] are computed for each cell and the values of these features at the two ends and at the center of each PC are computed for the shared and modality-specific latent spaces. The changes of representative morphological features along PC1 and PC16 of the chromatin shared latent space are plotted. (b) Heatmaps show the proportion of times each representative morphological feature is explained by a top PC in the shared or the modality-specific latent space. (c) Heatmap shows the morphological feature values in each cell, where each row represents a cell. The bar plot shows the fold change of prediction accuracy of each feature ablation compared to the prediction accuracy of using all features.

To more comprehensively characterize the information captured by each latent space, we compare the image features contained in the top PCs that explain 70% of total variance in each latent space (Figure 4b and SI Appendix, Figures S14c, S15c, S16c, and S19c). Interestingly, such analysis shows that heterochromatin volume, a feature associated with aging [38] and Alzheimer’s disease [15], is exclusively captured by the chromatin and protein shared latent space (namely in PC17, see Figure 4a). A similar analysis applied to each protein shows that foci count of *γ*H2AX is a protein-specific feature that is not captured by the chromatin and protein shared latent space (Figure 4b). When training separate regression models to predict the three *γ*H2AX features shown in Figure 4b based on chromatin images (SI Appendix, Extended Methods), the regression outputs show the lowest correlation with the ground truth feature values for the foci count of *γ*H2AX compared to the other two features that are mainly captured by the shared latent space (SI Appendix, Figure S18c). This indicates that APOLLO correctly identified *γ*H2AX foci count as a protein-specific feature while the other two features are also captured by chromatin imaging. A feature ablation test indicates that removing *γ*H2AX foci count results in the largest reduction of phenotype classification accuracy (Figure 4c) and is consistent with the significant reduction of phenotype classification accuracy observed when the protein-specific latent space of *γ*H2AX is not used in reconstructing the protein images (Figure 3d). This confirms that the modality-specific latent space can capture important disease-relevant features and demonstrates how APOLLO can be used to interrogate the shared and modality-specific information between imaging modalities, where shared features are not directly available.

### 6 APOLLO identifies novel associations between protein subcellular localization and the morphology of cellular components

Function and activity of a protein is known to be tightly coupled to its subcellular localization, which can vary across single cells even within the same cell line [27, 39, 40, 35, 41]. Computational tools have been developed to analyze single-cell variability in protein localization, showing that cellular and nuclear morphology can be used to predict protein subcellular localization in single cells [42, 36, 43]. However, these models take as input images of multiple cellular components and little is known about the association of each cellular component with the change in protein localization across single cells. We apply APOLLO to images of U2OS cells (the most abundant cell line) in the Human Protein Atlas (HPA) [27] to learn shared and modality-specific information between the morphology of nucleus, microtubule, and ER with respect to variations in protein subcellular localization (SI Appendix, Extended Methods). In HPA, all cells are stained for nucleus, microtubule and ER (the three reference stains), together with one additional target protein (total of 11657 proteins across all U2OS cells, see SI Appendix, Extended Methods). Concentrating on the 25 proteins with the most variable subcellular localization (SI Appendix, Extended Methods), we disentangle the information in the three cellular components to analyze their association with the variability in protein subcellular localization.

Towards this, we separately train three APOLLO models (SI Appendix, Extended Methods) to learn the shared and modality-specific information between each pair of the three reference stains. We then cluster the cells in each latent space into two clusters and test for each of the 25 proteins whether the two clusters capture differences in their subcellular localization as measured by the proportion of protein localized within the cell nucleus (Figure 5a, SI Appendix, Extended Methods). Notably, the three models consistently indicate that variation in intra-nuclear localization of many proteins is differentially captured by the three different compartments. For example, morphological features of both ER and microtubule, but not the nucleus, are associated with the variability in intra-nuclear localization of DDB1 (Figure 5a-c), which is known to change localization in response to UV exposure [44]. Both the microtubule and ER modality-specific latent spaces indicate that cells with smaller cytoplasmic volume and lower intensity of microtubule and ER are associated with high intra-nuclear localization of DDB1 (Figure 5b-c). In contrast, the intra-nuclear localization of CLNS1A and C8orf59 is only associated with morphological features of the nucleus (Figure 5a,d-e). Interestingly, for CLNS1A, a protein involved in small nuclear ribonucleoprotein biogenesis and control of cell volume [45, 46, 47], our model suggests that higher intra-nuclear localization is found in cells with higher heterochromatin content (Figure 5d); for C8orf59, a protein involved in ribosome biogenesis [48], our model suggests that increased intra-nuclear localization is associated with more circular nuclei (Figure 5e).

**Figure 5.**
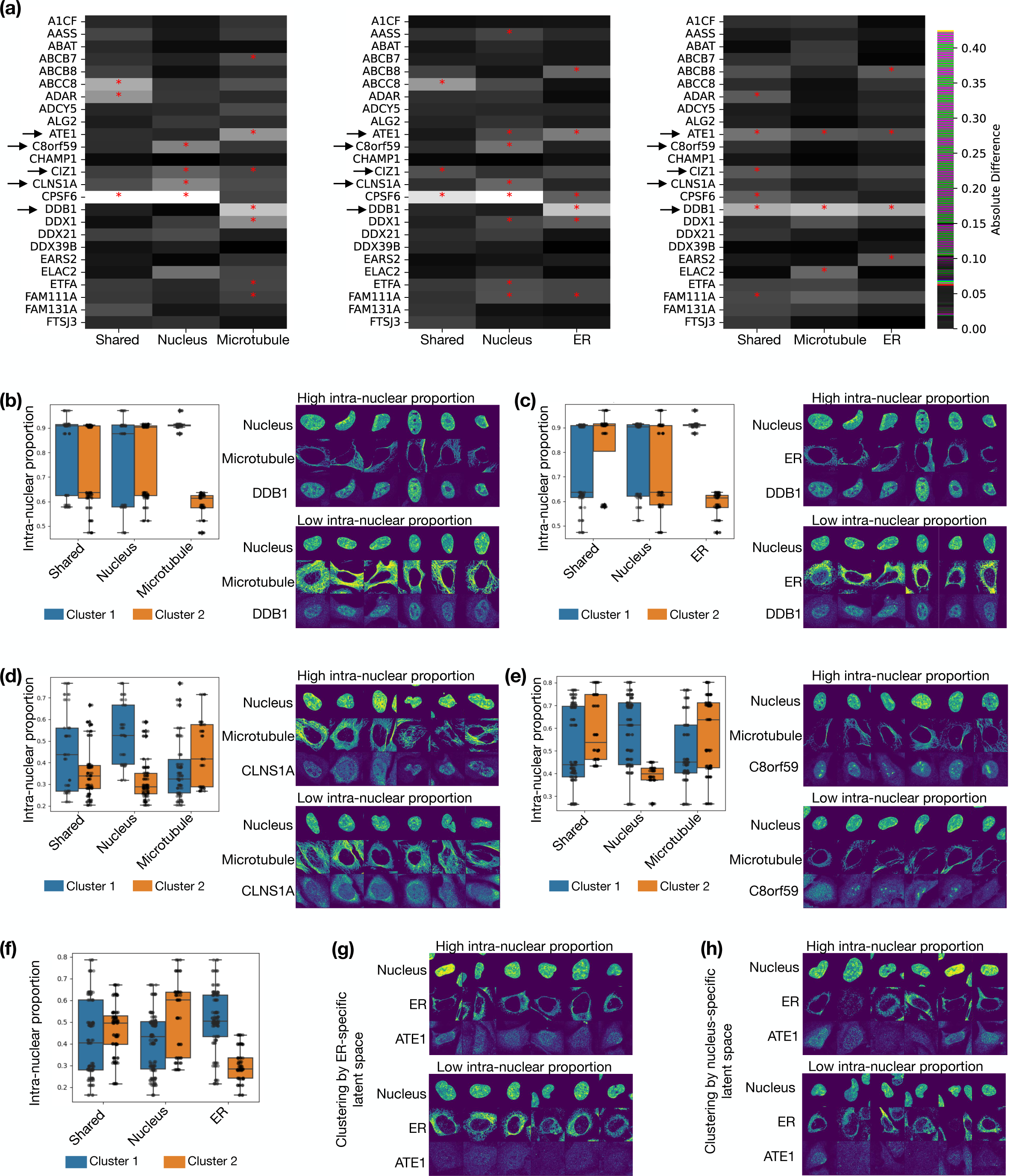
APOLLO disentangles the information contained in different cellular components with respect to protein subcellular localization. (a) We train three separate APOLLO models using nucleus and microtubule stains (left), nucleus and ER stains (middle), and microtubule and ER stains (right). For each model, each of the shared and the two modality-specific latent spaces is clustered into two clusters. The heatmap shows the absolute value of the difference in the mean intra-nuclear proportion in each cluster, averaged across the 5 random initializations (SI Appendix, Extended Methods). *: p-value *<* 0.00022 using two-sided t-test and absolute difference between clusters *>* 0.1 in all 5 random initializations (SI Appendix, Extended Methods). Arrows indicate proteins with example images shown in Figure 5 and SI Appendix, Figure S20. (b)-(e) Left: Box plot of the intra-nuclear proportions of DDB1 (b)-(c), CLNS1A (c), and C8orf59 (d) in the two clusters of each latent space, obtained using the models trained with either the nucleus and microtubule images (b, d, e) or the nucleus and ER images (c). Right: Examples of cells in each cluster of the microtubule-specific latent space (b), the ER-specific latent space (c) or the nucleus-specific latent space (d, e) stained for the particular proteins. (f) Box plot of the intra-nuclear proportions of ATE1 in the two clusters of each latent space, obtained using the model trained with nucleus and ER images. (g) Examples of cells stained for ATE1 in each cluster of the ER-specific latent space. (h) Examples of cells stained for ATE1 in each cluster of the nucleus-specific latent space.

For some proteins, the difference in intra-nuclear localization is captured by more than one modality-specific latent space but not the shared space in the same model; for example, both the nucleus-specific and ER-specific latent spaces but not the shared space capture differences in intra-nuclear localization of ATE1 across cells (Figure 5a). This indicates that multiple aspects of nuclear and cellular morphology, captured by the different latent spaces, could simultaneously contribute to the localization of a particular protein in different ways. Indeed, analyzing the clusters in the different modality-specific latent spaces shows that while they are associated with differences in the intra-nuclear localization of a protein, the cell clusters are different (SI Appendix, Figure S20a-c, Extended Methods). For example, while both the ER-specific and nucleus-specific latent spaces show significant separation of cells by their intra-nuclear localization of ATE1 (Figure 5a,f), visualizing cells in the clusters of the different latent spaces suggests that the ER-specific clusters capture differences in ER intensity and cytoplasmic volume (Figure 5g), while the nucleus-specific clusters capture differences in nuclear sizes (Figure 5h). A similar analysis for CIZ1 of the clusters in the microtubule-specific latent space suggests that low intra-nuclear localization is associated with larger ratio of cytoplasmic-to-nuclear volume (SI Appendix, Figure S20e), which is not a distinctive feature between the clusters of the nucleus-specific latent space (SI Appendix, Figure S20d). This analysis demonstrates that APOLLO can generalize to different multiplexed imaging experiments and provide novel insights on relations between protein subcellular localization and the morphology of different cellular components.

## Discussion

Recent advances in experimental techniques have enabled the simultaneous measurement of multiple modalities, including multiplexed imaging of proteins at the sub-cellular level [1, 27], SHARE-seq in the single-cell domain, and various spatial transcriptomic methods in the tissue context [7, 8, 9, 5]. Taking spatial transcriptomics in the context of Alzheimer’s disease as an example, we demonstrated in earlier work the importance of integrating information contained in chromatin and plaque staining, gene expression, and cellular neighborhoods to identify mouse cortex regions that are at different stages of disease progression [15]. In addition to incorporating multiple modalities to comprehensively characterize cell states, obtaining insights into the regulatory relationship between cellular components also requires understanding the difference in information captured between modalities. Thus, methods are needed that can integrate different modalities and at the same time tease apart information that is not shared between the different modalities.

We introduced a general computational framework for the analysis of multi-modal data, which explicitly models the shared and modality-specific information through the use of partially overlapping latent spaces. Our model is applicable to any paired modalities and does not require their alignment through a common coordinate system such as genes for paired sequencing data. While learning partially overlapping latent spaces is a difficult optimization problem, which cannot be solved through standard autoencoder training, we demonstrated in four different applications that our proposed training procedure using latent optimization is able to learn these novel and intricate latent space structures. In addition, we provided a systematic approach to associate interpretable features with the shared and modality-specific latent spaces.

When applied to paired scRNA-seq and scATAC-seq data from SHARE-seq of mouse skin cells [5], we found that cell type classification is more accurate when adding either the RNA-specific latent space or the ATAC-specific latent space compared to using the shared latent space alone, indicating that biologically meaningful information is captured in each of the latent spaces. Traversing the principal components of the shared and modality-specific latent spaces allows their interpretation by identifying the gene programs captured in the different latent spaces, without the need for manually annotating and comparing the clusters in each modality. To demonstrate the general applicability of APOLLO, we also applied APOLLO to paired scRNA-seq and cellular surface protein abundance data measured using CITE-seq [17]. Clustering each modality separately indicates that both modalities contain cell type information while only scRNA-seq contains experimental batch differences (SI Appendix, Figure S3 b-c). APOLLO correctly disentangles the shared and modality-specific information by capturing the cell-type difference in the shared latent space and the batch difference only in the RNA-specific latent space.

APOLLO can also be applied to more challenging datasets where the different modalities cannot directly be summarized using common features (such as genes in SHARE-seq or CITE-seq). We demonstrated this in the context of multiplexed imaging to study the association between chromatin organization and protein localization, using a dataset of PBMCs from different tumor patients [34]. Importantly, interpreting the shared and modality-specific latent spaces allows analyzing which aspects of chromatin organization and protein localization are shared and thus characterize the same aspects of the underlying cell state, as well as identifying protein localization features that are not captured by chromatin organization, indicating the presence of other regulatory mechanisms. APOLLO also allows for cross-modality prediction of protein localization based on chromatin images, which is a challenging task with limited previous work [36]. We demonstrated that our generated protein images on held-out patients outperforms previous methods and are similarly predictive of cancer type as the real protein images. Furthermore, our approach for interpreting the latent spaces provides means to understand what information is not captured by chromatin organization, thereby allowing a more informed assessment of applications using the predicted protein images.

In addition to learning the association between protein localization and chromatin organization, as shown using the PBMC data, we also demonstrated that APOLLO can incorporate additional reference stains to disentangle the information contained in the morphology of different cellular components with respect to protein localization. By clustering cells in the shared and modality-specific latent spaces of three reference stains (nucleus, ER, and microtubule), we examined which latent spaces capture difference in intra-nuclear protein localization across cells. In this way, we were able to associate variability in protein subcellular localization with morphological changes in the different cellular components.

Collectively, APOLLO enables explicit learning of shared and modality-specific information leading to a more holistic understanding of cell state and the underlying regulatory mechanisms. While we demonstrated APOLLO on SHARE-seq, CITE-seq and multiplexed protein staining, by adjusting the encoder and decoder structures, our method is broadly applicable to any multi-modal measurements to study cell state changes and regulatory mechanisms underlying development, disease, and other biological processes from single-cell data as well as well beyond the single-cell domain. For example, large-scale biobanks capturing paired information on the genetics and different medical modalities of individuals are being collected [23, 24]. APOLLO offers a framework to go beyond multi-modal integration [49] to disentangle the information contained in different medical modalities as well as to directly disentangle and associate genetic variants with particular variables of a medical measurement or health history.

## Methods

All model architectures, hyperparameters, and procedures for model training, evaluation, and interpretation used in the four applications are described in SI Appendix, Extended Methods. All datasets used are publicly available [5, 17, 34, 27], and pre-processing of the data was performed following standard procedures, which are described in detail in SI Appendix, Extended Methods.

## Acknowledgments

We thank all members of the Uhler and Shivashankar labs for discussions and E. Forte for editing the manuscript. X.Z. was supported by the Eric and Wendy Schmidt Center at the Broad Institute. GVS was partially supported by the Swiss National Foundation (310030 208046). CU was partially supported by NCCIH/NIH (1DP2AT012345), ONR (N00014-22-1-2116), AstraZeneca, the MIT-IBM Watson AI Lab, MIT J-Clinic for Machine Learning and Health, and a Simons Investigator Award.

## Code availability

The code is available in the Github repository: https://github.com/uhlerlab/APOLLO/.

