## Supplementary Information for "Partially Shared Multi-Modal Embedding Learns Holistic Representation of Cell State"

**This PDF file includes:**
Extended Methods
Figures S1-S20
SI References

### Extended Methods

#### 1 Model architecture and training of paired scRNA-seq and scATAC-seq

For both training steps, we split the data into training and validation sets, as is standard in neural network training. We use the same randomly selected 85% of cells from the SHARE-seq dataset [10] to train our model and the remaining 15% to validate and test the generalization performance of our model.

##### 1.1 Step 1 - latent optimization

In this step, we train the latent spaces and the corresponding decoders to reconstruct the scRNA-seq and scATAC-seq data (SI Appendix, Figures S1b and S2a).

**Latent spaces of paired scRNA-seq and scATAC-seq:** The shared latent space has 50 dimensions and each of the two modality-specific latent spaces have 20 dimensions. The shared latent space is chosen to be much larger than the modality-specific latent spaces to ensure that the shared space has enough capacity to contain all shared information. Latent spaces are initialized using "torch.nn.Embedding" with the default parameters. Independent Gaussian noise with zero mean and unit variance is added to the latent spaces at each training epoch. The hyperparameter $\lambda$ in Equation (1) for the $\ell_2$ regularization of the latent spaces is set to 0.001 and the learning rate of the ADAM optimizer is set to 0.001, which are the same hyperparameter values as in [3].

**Decoders of paired sequencing-based modalities:** Four decoders are trained in step 1: a) decoder that reconstructs scRNA-seq from the shared latent space; b) decoder that reconstructs scRNA-seq from the full RNA-seq latent space, i.e., the shared latent space embedding concatenated with the RNA-specific latent space embedding; c) decoder that reconstructs scATAC-seq from the shared latent space; d) decoder that reconstructs scATAC-seq from the full ATAC-seq latent space, i.e., the shared latent space embedding concatenated with the ATAC-specific latent space embedding. The architectures of the decoders are shown in Figure S1b. The input feature dimension is 70 for the full latent space decoder and 50 for the shared latent space decoder except when different latent space sizes are tested (Figure 2a). Each decoder has 3 hidden layers, each of which has dimension 1024. All hidden layers are linear layers with a dropout rate of 0.01 and are followed by LeakyReLU activation and a batch normalization layer. The output layer is a linear layer with a dropout rate of 0.01 and sigmoid activation. This decoder architecture follows standard autoencoder setups [8, 15]. For both modalities, we use binary cross-entropy loss to minimize the objective function defined in Equation (1), which is calculated using "torch.nn.BCEWithLogitsLoss" based on the output before the sigmoid activation with pos_weight set inversely proportional to the total counts of genes or peaks. The learning rate of the decoders is 0.0001 and ADAM is used for optimization, which is the same as in [3].

#### 1.2 Step 2 - inference

In this step, we train the encoders to infer the learned latent spaces from the scRNA-seq and scATAC-seq data without updating the latent spaces or the decoders (SI Appendix, Figures S1c and S2b).

**Encoders of paired scRNA-seq and scATAC-seq:** We train two separate encoders for scRNA-seq and scATAC-seq with identical structures (SI Appendix, Figure S1c). Each encoder starts with two linear layers with a dropout rate of 0.01, each of which has 1024 dimensions and is followed by LeakyReLU activation and batch normalization, which is a standard setup for autoencoders [8, 15]. After the first two hidden layers, two separate linear layers with LeakyReLU activation are used to obtain separate hidden layers for the shared and modality-specific latent spaces. A linear layer is applied to each hidden layer to obtain the shared or modality-specific latent space. The inferred latent spaces from the autoencoder are compared to the latent spaces learned in step 1 through mean-squared error (MSE) loss. The MSE loss is minimized using the ADAM optimizer with a learning rate of 0.0001, the same learning rate as the decoder.

#### 1.3 Application to paired scRNA-seq and protein abundance

The same setup is applied to the paired scRNA-seq and protein abundance data measured by CITE-seq [4], with dimension 50 for the shared latent space and dimension 30 for the modality-specific latent space. Given that cross-modality prediction isn't the goal in this application and to demonstrate robustness of our approach, the model is trained without the two decoders $D_1'$ and $D_2'$ that map from the shared latent space to each of the modalities (SI Appendix, Figure S3a).

### 2 Pre-processing of scRNA-seq and scATAC-seq data

The paired scRNA-seq and scATAC-seq datasets from [10] are separately filtered for genes or peaks that are non-zero in at least 300 cells and then filtered for cells with at least 300 non-zero genes or peaks. The common cells between the two modalities are then selected for another round of filtering with the same criteria. These two rounds of filtering result in 28,098 cells with 9,153 genes in the scRNA-seq data, and 58,170 peaks in the scATAC-seq data. The data is then log-transformed and min-max scaled, such that the minimum count in each cell is 0 and the maximum count in each cell is 1. Our filtering and normalization steps follow standard procedures of pre-processing scRNA-seq and scATAC-seq data; see e.g. [10, 14, 2, 15].

### 3 Preprocessing of scRNA-seq and cellular surface protein abundance data

Following the approach taken by a previous study that analyzes this dataset [4], we use the top 4005 highly variable genes in the scRNA-seq data and all 110 proteins. Same as in the pre-processing of scRNA-seq and scATAC-seq described above, the data is log-transformed and min-max scaled, such that the minimum count in each cell is 0 and the maximum count in each cell is 1.

#### 4 Alternative models for scRNA-seq and cellular surface protein abundance data

We use the default parameter setting for the Seurat WNN method [6]. For testing the standard multi-modal autoencoder method, we use the same encoder and decoder architectures as the full APOLLO model with 80 latent dimensions, which is equal to the full latent space dimension of the APOLLO model. Instead of performing a two-step training, this model trains the encoder and decoder together without directly updating the latent spaces. Mean-squared error loss is added between the latent spaces obtained from the two encoders of the two input modalities. Similar to the APOLLO model that adds Gaussian noise in the latent space, this model performs a variational sampling step as in a standard variational autoencoder and the resulting latent spaces are passed to the decoder for reconstruction. In each epoch, we alternate between training the scRNA-seq autoencoder and training the protein autoencoder.

#### 5 Cell type prediction based on the inferred latent spaces of scRNA-seq and scATAC-seq

We train four separate neural network classifiers with the same architecture to predict cell types based on one of the following inputs: a) shared latent space inferred from scRNA-seq data using the RNA encoder; b) shared latent space inferred from scATAC-seq using the ATAC encoder; c) both shared and modality-specific latent spaces inferred from scRNA-seq data using the RNA encoder; d) both shared and modality-specific latent spaces inferred from scATAC-seq data using the ATAC encoder. We use a standard feedforward neural network architecture in our classifiers; see e.g. [5]. Each classifier consists of four layers and outputs the probability of a given cell being assigned to each of the 23 cell types. The first three layers are followed by LeakyReLU activation and batch normalization. All four layers have a dropout rate of 0.1 and a hidden dimension of 128. We use the ADAM optimizer with a learning rate of 0.00001 to minimize the cross entropy loss between our prediction and the cell type labels assigned by Ma *et al.* [10]. We use the same train-validation split to train the classifiers as in the training of the APOLLO model, i.e., the same 85% of cells are used to train the APOLLO model and the cell type classifiers.

#### 6 Model architecture and training of paired chromatin and protein imaging

For both training steps, we hold-out all images from one patient in each of the four phenotypes for testing the model (SI Appendix, Figure S6).

##### 6.1 Step 1 - latent optimization

In this step, we train the latent spaces, protein IDs, and the corresponding decoders to reconstruct the chromatin and protein images (SI Appendix, Figures S7a and S8a).

**Latent spaces and protein IDs of paired chromatin and protein imaging:** For each protein, we randomly initialize a trainable 64-dimensional vector of protein ID that is shared across all images of that protein by using "torch.nn.Embedding". The protein IDs are used in both the encoding and decoding steps by concatenating them to the latent space embeddings, similar to a conditional autoencoder model (SI Appendix, Figure S7). The shared latent space has 1024 dimensions and each of the two modality-specific latent spaces has 200 dimensions except when different latent space sizes are tested (Figure 3b). Similar to the application to paired scRNA-seq and scATAC-seq, the shared latent space is chosen to be much larger than the modality-specific latent spaces to ensure that the shared space has enough capacity to contain all the shared information. Latent spaces are initialized using "torch.nn.Embedding" with the default parameters. Independent Gaussian noise with zero mean and unit variance is added to the latent spaces at each training epoch. The hyperparameter $\lambda$ in Equation (1) for the $\ell_2$ regularization of the latent spaces is set to 0.001 and the learning rate of the ADAM optimizer is set to 0.001, which are the same hyperparameter values as in [3].

**Decoders of paired chromatin and protein imaging:** Four decoders are trained in step 1: a) decoder that reconstructs chromatin images from the shared latent space; b) decoder that reconstructs chromatin images from the full chromatin latent space, i.e., the shared latent space embedding concatenated with the chromatin-specific latent space embedding; c) decoder that reconstructs protein images from the shared latent space; d) decoder that reconstructs protein images from the full protein latent space, i.e., the shared latent space embedding concatenated with the protein-specific latent space embedding. The architectures of the decoders are shown in SI Appendix, Figure S7a. The latent space embedding, after adding noise and being concatenated with the protein ID, is passed through a linear layer with ReLU activation and reshaped to $4 \times 4 \times 96$ dimensions for subsequent convolutions. This is followed by 5 convolutional layers with a kernel size of 4 and stride of 2. The number of channels in each hidden layer is listed in SI Appendix, Figure S7a. The first four convolutional layers are followed by batch normalization and LeakyReLU activation. The last convolutional layer is followed by sigmoid activation to scale the output image from 0 to 1. For both imaging modalities, we use binary cross-entropy loss to minimize the objective function defined in Equation (1), which is calculated using "torch.nn.BCEWithLogitsLoss" based on the output before the sigmoid activation. The learning rate of the decoders is 0.0001 and ADAM is used for optimization. The decoder architectures and training procedure follow standard setups for autoencoders; see e.g. [3, 9, 15].

#### 6.2 Step 2 - inference

In this step, we train the encoders to infer the learned latent spaces from the chromatin and protein images without updating the latent spaces, protein IDs, or the decoders (SI Appendix, Figures S7b and S8b).

**Encoders of paired chromatin and protein imaging:** We train two separate encoders for chromatin and protein images with identical structures (SI Appendix, Figure S7b). Each encoder starts with 5 convolutional layers with LeakyReLU activation, a standard setup for autoencoders [3, 9, 15]. The dimensions of the hidden layers are listed in SI Appendix, Figure S7b. The output of the last convolutional layer is divided into two sets of channels that are used to derive the shared and modality-specific latent spaces respectively. 80 out of the 96 channels of the last hidden layer are flattened, concatenated with protein ID, and passed through a linear layer to obtain the shared latent space. Similarly, the modality-specific latent space is obtained from the remaining 16 channels. The inferred latent spaces from the autoencoder are compared to the latent spaces learned in step 1 through mean-squared error (MSE) loss. The MSE loss is minimized using the ADAM optimizer with a learning rate of 0.001.

#### 6.3 Cross-modality predictions

To predict unmeasured proteins, the shared latent space is first inferred from the chromatin image of a cell using the chromatin encoder. Then each protein image can be predicted by decoding the inferred shared latent space through the protein decoder using the protein ID of the target protein (SI Appendix, Figure S7a)

#### 6.4 Application to Human Protein Atlas data

The same setup is applied to the Human Protein Atlas data [12], with a shared latent space dimension of 1024 and a modality-specific latent space dimension of 200. The two decoders $D'_1$ and $D'_2$ that map from the shared latent space to each of the modalities are not used, since the omission does not impact cross-modality-prediction performance (Figure 3b) and results in correct disentanglement of the shared and modality-specific information (Figure 2f).

#### 6.5 Alternative models

The models used for benchmarking our full APOLLO model are described below. In addition to using BCE loss for the decoder outputs, we also tested the use of an alternative loss function, namely the mean-squared error (MSE) loss. All model training and testing use the same train-test split. For benchmarking cross-modality prediction, the prediction results are first thresholded using mode intensity and then compared using $\ell_1$ loss, which is not used for training any of the models.

**Standard autoencoder training using a single step (SI Appendix, Figure S12a):** This model has the same encoder and decoder architectures as the full APOLLO model, and the same dimensions of the shared, chromatin-specific, and protein-specific latent spaces are used. Protein IDs are concatenated to the hidden layers during encoding and decoding as described in the full model. However, instead of performing a two-step training, this model trains the encoder and decoder together without directly updating the latent spaces. Mean-squared error loss is added between the shared latent spaces obtained from the two encoders of the two input modalities. Similar to the APOLLO model that adds Gaussian noise in the latent space, this model performs a variational sampling step as in a standard variational autoencoder and the resulting latent spaces are passed to the decoder for reconstruction. In each epoch, we alternate between training the chromatin autoencoder and training the protein autoencoder.

**Our model without modality-specific latent space (SI Appendix, Figure S12b):** This model has the same two-step training procedure as the full APOLLO model, but does not separate the shared and the modality-specific latent spaces. There are only two decoders that decode the two modalities from the shared latent space and the are trained in step 1. The encoders only output the shared latent space, which has the same dimension as the combined dimension of the shared and the modality-specific latent spaces in the full APOLLO model.

**Inpainting model developed by Lu *et al.* [9]:** We adapted the original code from TensorFlow to PyTorch. We used the same hyperparameter values as in the original publication, including the use of mean-squared error loss, except the slight modification to adapt to the single-channel input image without a microtubule channel.

#### 7 Pre-processing of paired chromatin and protein imaging

The imaging data is obtained from [1], where the patients are annotated with one of the following four phenotypic classes: healthy, meningioma, glioma, or head and neck tumor (SI Appendix, Figure S6). Each single-cell image of each protein stain and chromatin stain is min-max scaled such that the minimum pixel value is 0 and the maximum is 1, a standard image normalization step for neural networks [1, 9]. For each cell nucleus an image patch centered at the centroid of the nucleus of dimension $128 \times 128$ pixels is cropped from the whole image.

#### 8 Pre-processing of the Human Protein Atlas images

The same procedure as in a previous study [16] is used to pre-process the images. We use all images of U2OS cells (the cell line with the most data in the Human Protein Atlas) that are also used in [16] for studying protein localization variabiltiy (2973 proteins used) and exclude proteins that are stained in less than 150 cells, resulting in a total of 141 proteins used for training the model. Nuclear segmentation obtained using StarDist [13] is used for computing the intra-nuclear proportion of the protein in each image. The standard deviation of the intra-nuclear proportion of each protein is computed across all images stained for the protein. The 25 proteins with the largest standard deviations are used for interpreting the disentanglement results (Figure 5).

#### 9 Phenotype classification using real or reconstructed protein and chromatin images

We use the ResNet-18 model [7], a popular neural network model for image classification, in PyTorch for the phenotype classifiers to predict phenotypes from real protein/chromatin images, reconstructed protein/chromatin images, or protein images predicted from chromatin (Figure 3d). The first layer of the ResNet-18 model is adjusted to take single-channel input images. We use the cross entropy loss and set the weight of each phenotype class to be inversely proportional to the fraction of cells in that particular phenotype class. The classifiers are trained using the ADAM optimizer with a learning rate of 0.001. The training of each classifier is repeated 36 times with 6 different groups of held-out patients, where each held-out group contains one patient from each phenotype class. One-sided t-test is used for the null hypothesis that the phenotype prediction accuracy using the reconstruction from the full latent space is smaller than or equal to the accuracy using the reconstruction from the shared latent space.

#### 10 Interpretation of the partially shared latent spaces

In the following, we describe a procedure to analyze the shared latent space as well as the modality-specific latent space, which can be applied to any data modality. We compute the principal components (PCs) of the latent spaces and group the cells based on their positions along the PCs. For both of our applications, we group cells into 11 bins with equal percentile ranges along each PC. In the following description of our approach, we denote the percentile bin of $PC_i$ that is centered around 0 as $PC_i^0$ and the bins ordered from negative to positive PC values are denoted as $PC_i^{-5}, ...PC_i^{-1}, PC_i^0, PC_i^1, ...PC_i^5$. When comparing cells along $PC_i$ we only use the cells at the center of all other PCs, i.e. the 15% of cells with the smallest Euclidean distance to the PC origin calculated using all $PC_j$ for $j \neq i$ and $j < 10$. The number of PCs considered in the analysis can be adjusted given the variance explained by each PC in a particular dataset. Using cells at the center of all other PCs, except the PC being interpreted, ensures that we are only considering the variation of cells along one PC to tease apart the variations explained by the different PCs. In the following, we explain this procedure in more detail in the context of the paired sequencing and paired imaging datasets, where we also perform subsampling for a more robust identification of the features explained by each PC.

##### 10.1 Interpretation of the latent spaces of paired scRNA-seq and scATAC-seq

We test for significant gene expression and peak count changes along each PC. The cells are divided into 36 random train-test splits with 20% of cells used for testing in each split, in order to validate the significant genes and peaks identified along each PC. We consider three groups of cells for each $PC_i$ that are all at the center of other PCs: 1) cells in $PC_i^{-5}$ and $PC_i^{-4}$; 2) cells in $PC_i^0$; 3) cells in $PC_i^4$ and $PC_i^5$. Cells in each group are compared to all cells in the other two groups through t-tests using Scanpy's "rank_genes_groups" function [14]. The threshold of p-values after multiple testing correction is set to 0.05 for both modalities. The number of times in all train-test splits a gene or peak is tested to be significant in both the training and testing cells with the same direction of fold change are plotted for different thresholds of log fold change magnitude (Figure 2c, and SI Appendix, Figures S4, and S5). The ATAC-seq peaks in the shared and modality-specific latent spaces are summarized in Figure 2d, respectively. For this, we counted the number of PCs each peak is found to be significant in the shared and modality-specific latent spaces respectively, grouped the genes by their gene ontology annotations, and normalized the total counts in the two latent spaces to 1.

##### 10.2 Interpretation of the latent spaces of paired chromatin and protein imaging

Imaging data can be visually inspected by sampling cells at each percentile bin along a particular PC (Figure 4a). Additionally, for a more comprehensive assessment, predefined handcrafted image features [11, 1] can be used to test for significant feature changes along each PC. For the following analysis, we use a total of 234 chromatin image features and around 30 image features per protein obtained from [1]. The patients are divided into 12 different train-test splits, which are summarized in SI Appendix, Figure S6, in order to validate the significant features identified along each PC. As in the application to paired scRNA-seq and scATAC-seq, we test for significant feature changes among three groups of cells for each $PC_i$ that are all at the center of other PCs: 1) cells in $PC_i^{-5}$ and $PC_i^{-4}$; 2) cells in $PC_i^0$; 3) cells in $PC_i^4$ and $PC_i^5$. A feature is considered significant if the p-value after multiple testing correction is less than 0.05 and the absolute value of log fold change is greater than $log_2(fc_t)$ in both the training and testing patients, where $fc_t$ is set to be 1.2 for chromatin features and 1.4 for protein features. The fold-change thresholds are chosen so that most representative morphological features are represented by at least one PC. Additionally, the direction of fold change is required to be the same in the training and testing patients. Figure 4a and SI Appendix, S14 shows the chromatin features that are significant in at least 8 out of the 12 train-test splits (SI Appendix, Figure S6). We summarize the morphological features in all top PCs in the shared and modality-specific latent spaces respectively in Figure 4b, and SI Appendix, Figures S14c, S15c, S16c, and S19c. For this, we counted the number of PCs each feature is found to be significant in the shared and modality-specific latent spaces and normalized the total counts in the two latent spaces to 1.

To obtain a more concise representation, we group the morphological features of chromatin and each protein by Pearson correlation across all cells and plot one representative feature per group in Figure 4a, 4b, and SI Appendix Figures S13-19. The feature groupings for each stain are obtained as follows: we build a network with the features as nodes and an edge is added between each pair of features whose Pearson correlation is greater than 0.7. Features within the same connected component of the graph are considered in the same group. The representative feature of a group is the node with the highest degree. This results in 18 feature groups for chromatin, 5 groups for $\gamma$H2AX, 5 groups for lamin, 6 groups for CD8, 4 groups for CD4, 5 groups for CD3, and 4 groups for CD16.

#### 10.3 Interpretation of the latent spaces of the Human Protein Atlas images

For each subset of U2OS cells stained with the same protein, we separately perform k-means (k = 2) clustering in the shared and the two modality-specific latent spaces of each model. The clustering in each latent space is repeated 5 times with different random seeds. For each random seed, t-test is performed to compare the proportion of intra-nuclear protein localization in the two clusters, and the absolute value of the difference between the cluster means is computed. A clustering shows significant difference of intra-nuclear proportions if p-value $< 0.00022$ and absolute difference between clusters $> 0.01$ in all 5 random initializations. The p-value threshold is chosen by applying a Bonferroni correction to the overall probability of Type 1 error at 0.05 for a total of 225 tests performed for 25 proteins in each of the three latent spaces of the three separate models.

### 11 $\gamma$H2AX feature prediction using chromatin images

We use the ResNet-18 model [7] in PyTorch to train separate regression models that predict the predefined $\gamma$H2AX features from chromatin images. Similar to the previous section, for each of the three $\gamma$H2AX features (Figure 4b), a regression model is trained 12 times with different train-test splits of the patients (SI Appendix, Figure S6). The outputs of each regression model on the held-out patients are then compared to the ground truth using Pearson's correlation (SI Appendix, Figure S18c).

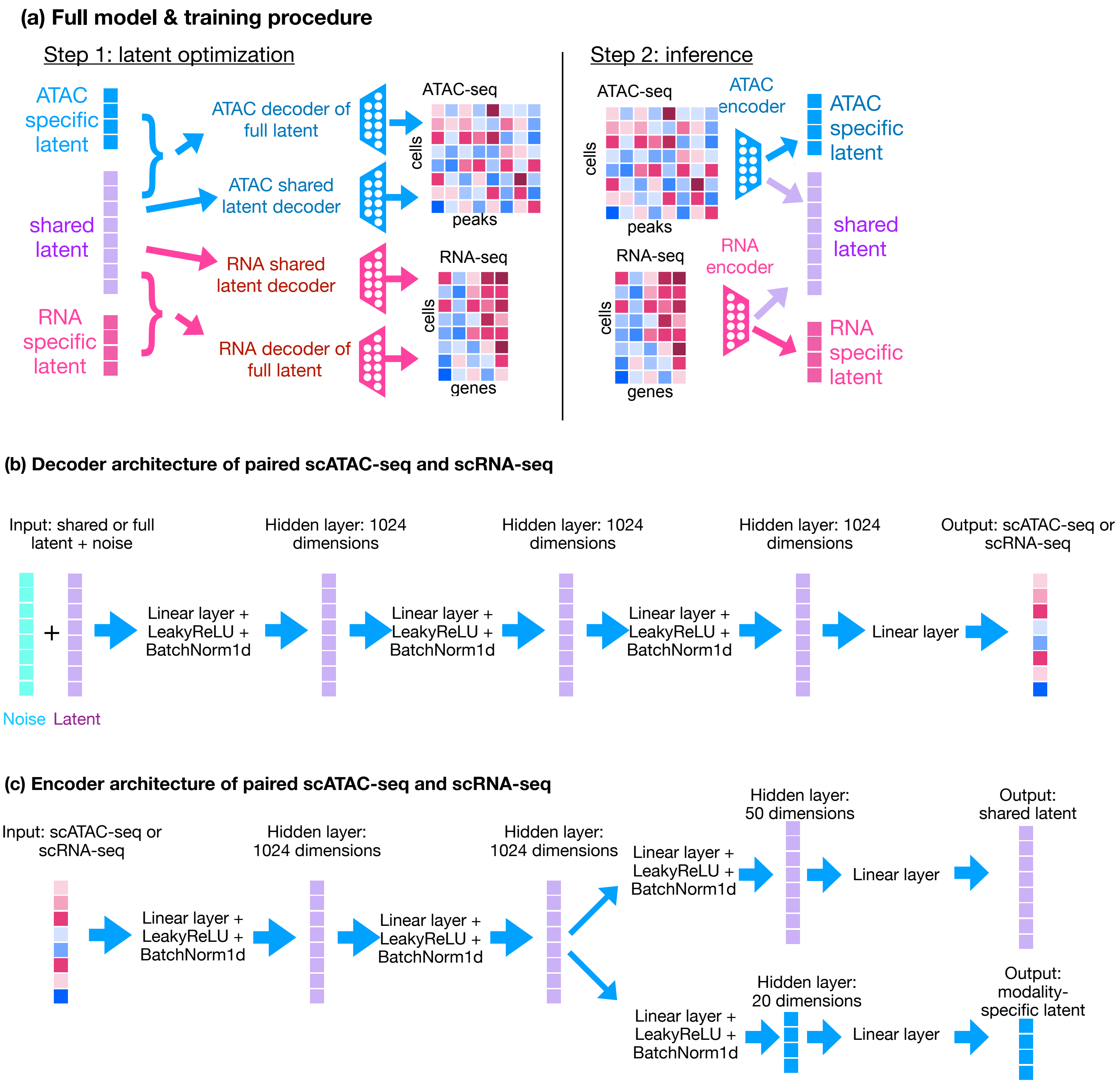

(a) Full model & training procedure
Step 1: latent optimization
ATAC specific latent
shared latent
RNA specific latent
ATAC decoder of full latent
ATAC shared latent decoder
RNA shared latent decoder
RNA decoder of full latent
ATAC-seq
cells
peaks
RNA-seq
cells
genes
Step 2: inference
ATAC-seq
cells
peaks
ATAC encoder
RNA-seq
cells
genes
RNA encoder
ATAC specific latent
shared latent
RNA specific latent
(b) Decoder architecture of paired scATAC-seq and scRNA-seq
Input: shared or full latent + noise
+
Noise
Latent
Linear layer + LeakyReLU + BatchNorm1d
Hidden layer: 1024 dimensions
Linear layer + LeakyReLU + BatchNorm1d
Hidden layer: 1024 dimensions
Linear layer + LeakyReLU + BatchNorm1d
Hidden layer: 1024 dimensions
Linear layer
Output: scATAC-seq or scRNA-seq
(c) Encoder architecture of paired scATAC-seq and scRNA-seq
Input: scATAC-seq or scRNA-seq
Linear layer + LeakyReLU + BatchNorm1d
Hidden layer: 1024 dimensions
Linear layer + LeakyReLU + BatchNorm1d
Hidden layer: 1024 dimensions
Linear layer + LeakyReLU + BatchNorm1d
Hidden layer: 50 dimensions
Linear layer
Output: shared latent
Linear layer + LeakyReLU + BatchNorm1d
Hidden layer: 20 dimensions
Linear layer
Output: modality-specific latent


**Figure S1**

**Figure S1. APOLLO architecture for paired scATAC-seq and scRNA-seq data.** (a) APOLLO uses a two-step training procedure. In step 1, the shared and modality-specific latent spaces as well as the modality-specific decoders are trained so that the decoders can reconstruct the scATAC-seq and scRNA-seq data from the latent spaces. For each modality, there is one decoder that reconstructs the modality using only the shared latent space and another decoder that reconstructs the modality using the full latent space, i.e. the shared latent space and the modality-specific latent space. In step 2, modality-specific encoders are trained to enable inferring the latent space embedding for cells not used in training the model and thus enable cross modality prediction. The encoders are trained to map the input features, e.g. scRNA-seq and scATAC-seq, to the latent space embeddings obtained in step 1.
(b) The decoders of scRNA-seq and scATAC-seq have the same architectures with four fully connected layers, except that the output layer decodes to different feature dimensions for the two modalities. The input feature dimension is 70 for the full latent space decoder and 50 for the shared latent space decoder. Each decoder has 3 hidden layers, each of which has dimension 1024. All hidden layers are linear layers with a dropout rate of 0.01 and are followed by LeakyReLU activation and a batch normalization layer. The output layer is a linear layer with a dropout rate of 0.01 and sigmoid activation.
(c) The encoders of scRNA-seq and scATAC-seq have the same architectures, except the difference in the input feature dimensions. Each encoder starts with two linear layers with a dropout rate of 0.01, each of which has 1024 dimensions and is followed by LeakyReLU activation and batch normalization, which is a standard setup for autoencoders [8, 15]. After the first two hidden layers, two separate linear layers with LeakyReLU activation are used to obtain separate hidden layers for the shared and modality-specific latent spaces. A linear layer is applied to each hidden layer to obtain the shared or modality-specific latent space.

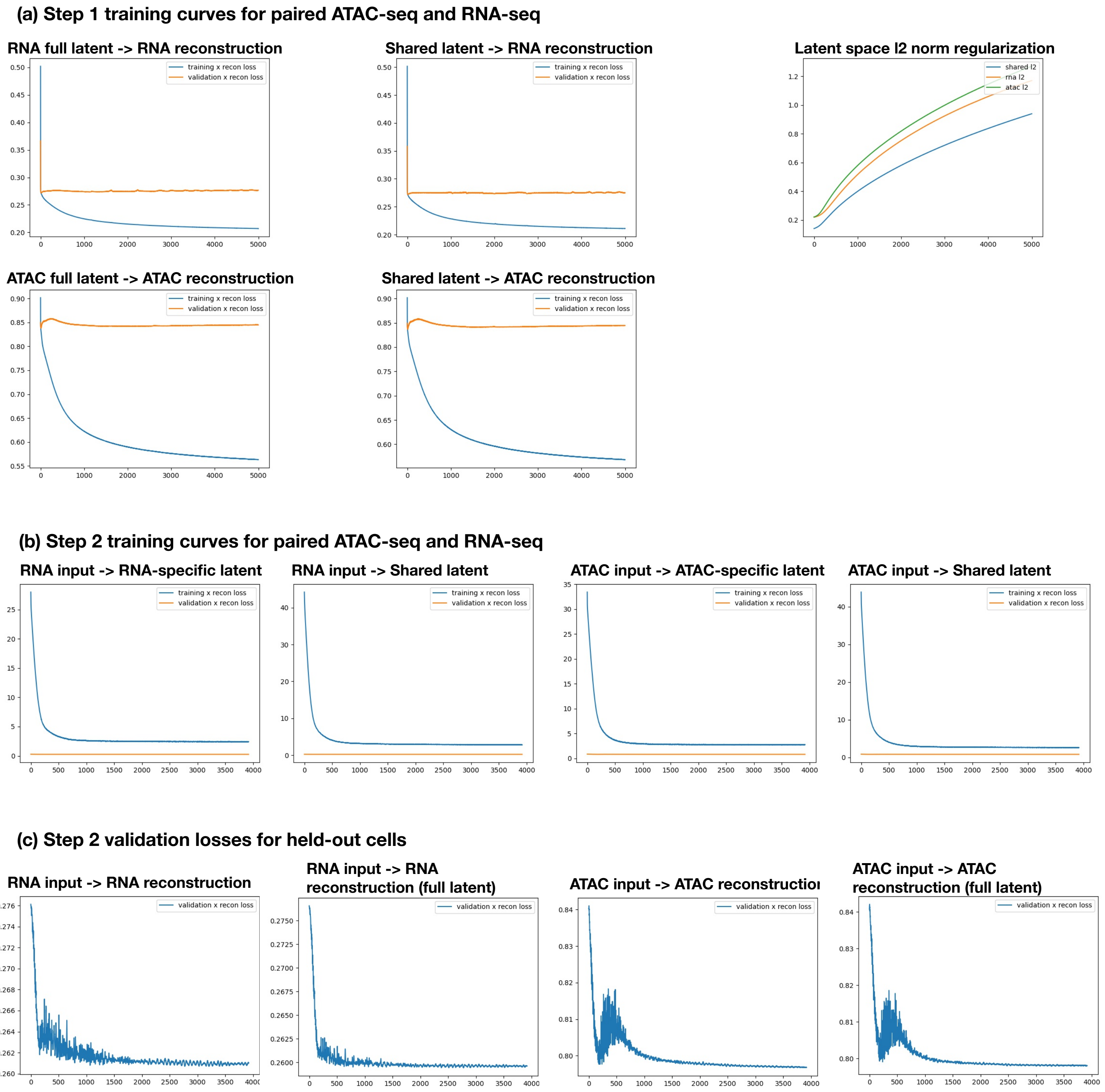
(a) Step 1 training curves for paired ATAC-seq and RNA-seq
RNA full latent -> RNA reconstruction
Shared latent -> RNA reconstruction
Latent space l2 norm regularization
shared l2
rna l2
atac l2
ATAC full latent -> ATAC reconstruction
Shared latent -> ATAC reconstruction
training x recon loss
validation x recon loss
(b) Step 2 training curves for paired ATAC-seq and RNA-seq
RNA input -> RNA-specific latent
RNA input -> Shared latent
ATAC input -> ATAC-specific latent
ATAC input -> Shared latent
(c) Step 2 validation losses for held-out cells
RNA input -> RNA reconstruction
RNA input -> RNA reconstruction (full latent)
ATAC input -> ATAC reconstructio
ATAC input -> ATAC reconstruction (full latent)

Figure S2

**Figure S2. Training and validation curves of APOLLO applied to paired scATAC-seq and scRNA-seq data.**
(a) Training curves are plotted for the latent optimization in step 1. Validation curves are expected to not decrease with training because the randomly initialized latent space of the validation samples are not updated during step 1 training. "RNA full latent → RNA reconstruction" shows the scRNA-seq reconstruction loss using the full RNA latent space, i.e., the shared latent space embedding concatenated with the RNA-specific latent space embedding. "Shared latent → RNA reconstruction" shows the scRNA-seq reconstruction loss using the shared latent space. "ATAC full latent → ATAC reconstruction" shows the scATAC-seq reconstruction loss using the full ATAC latent space, i.e., the shared latent space embedding concatenated with the ATAC-specific latent space embedding. "Shared latent → ATAC reconstruction" shows the scATAC-seq reconstruction loss using the shared latent space. "Latent space l2 norm regularization" shows the $\ell_2$ norms of the three latent spaces during training.
(b) Training curves for the inference in step 2 are plotted for the reconstruction of the shared and modality-specific latent spaces learned in step 1. "RNA input → RNA-specific latent" considers the reconstruction of the RNA-specific latent space from scRNA-seq data using the RNA-encoder. "RNA input → Shared latent" considers the reconstruction of the shared latent space from scRNA-seq data using the RNA-encoder. "ATAC input → ATAC-specific latent" considers the reconstruction of the ATAC-specific latent space from scATAC-seq data using the ATAC-encoder. "ATAC input → Shared latent" considers the reconstruction of the shared latent space from scATAC-seq data using the ATAC-encoder.
(c) RNA and ATAC reconstruction losses are plotted for cells held out from training. "RNA input → RNA reconstruction" shows the reconstruction loss of held-out cells obtained from first inferring the shared latent space using the RNA encoder and then decoding the scRNA-seq data using the RNA decoder of the shared latent space. "RNA input → RNA reconstruction (full latent)" shows the reconstruction loss of held-out cells obtained from first inferring the shared latent space and the RNA-specific latent space using the RNA encoder and then decoding the scRNA-seq data using the RNA decoder of the full RNA latent space. "ATAC input → ATAC reconstruction" shows the reconstruction loss of held-out cells obtained from first inferring the shared latent space using the ATAC encoder and then decoding the scATAC-seq data using the ATAC decoder of the shared latent space. "ATAC input → ATAC reconstruction (full latent)" shows the reconstruction loss of held-out cells obtained from first inferring the shared latent space and the ATAC-specific latent space using the ATAC encoder and then decoding the scATAC-seq data using the ATAC decoder of the full ATAC latent space.

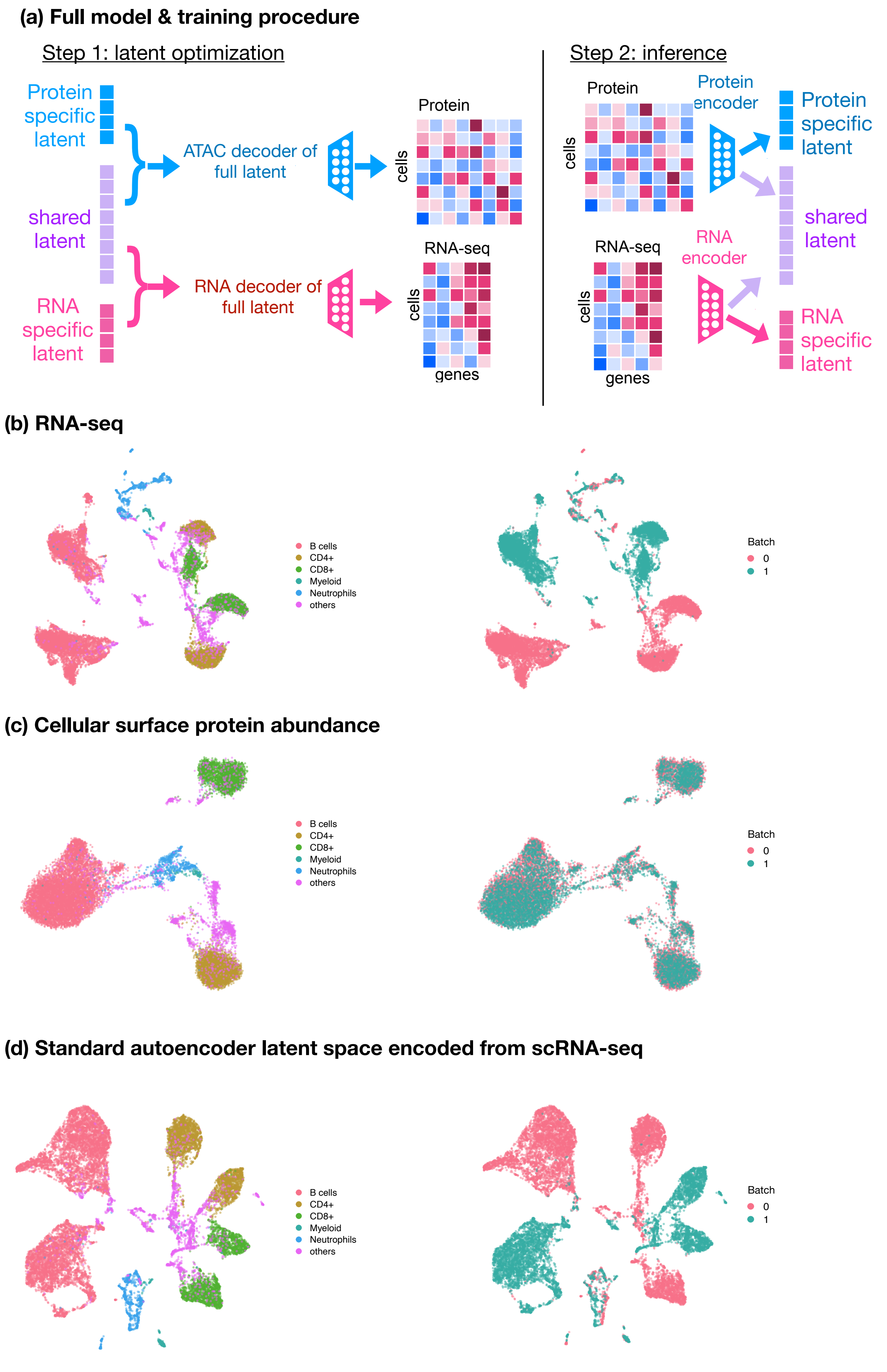

(a) Full model & training procedure
Step 1: latent optimization
Protein specific latent
shared latent
RNA specific latent
ATAC decoder of full latent
RNA decoder of full latent
Protein
cells
RNA-seq
cells
genes
Step 2: inference
Protein
cells
Protein encoder
Protein specific latent
shared latent
RNA-seq
cells
genes
RNA encoder
RNA specific latent
(b) RNA-seq
B cells
CD4+
CD8+
Myeloid
Neutrophils
others
Batch
0
1
(c) Cellular surface protein abundance
B cells
CD4+
CD8+
Myeloid
Neutrophils
others
Batch
0
1
(d) Standard autoencoder latent space encoded from scRNA-seq
B cells
CD4+
CD8+
Myeloid
Neutrophils
others
Batch
0
1


**Figure S3**

**Figure S3. Application of APOLLO to paired scRNA-seq and protein abundance data.**
(a) We applied the same encoder and decoder architecture as well as the default two-step training procedure to the paired scRNA-seq and protein abundance data (Extended Methods).
(b) UMAP visualization of scRNA-seq data using the Scanpy package [14].
(c) UMAP visualization of the protein abundance data using the Scanpy package [14].
(d) UMAPs obtained from the shared latent space of standard autoencoders, encoded using scRNA-seq data, separately colored by cell types and experimental batches.

**(a) Top genes explained by principal components of scATAC-seq shared latent space**

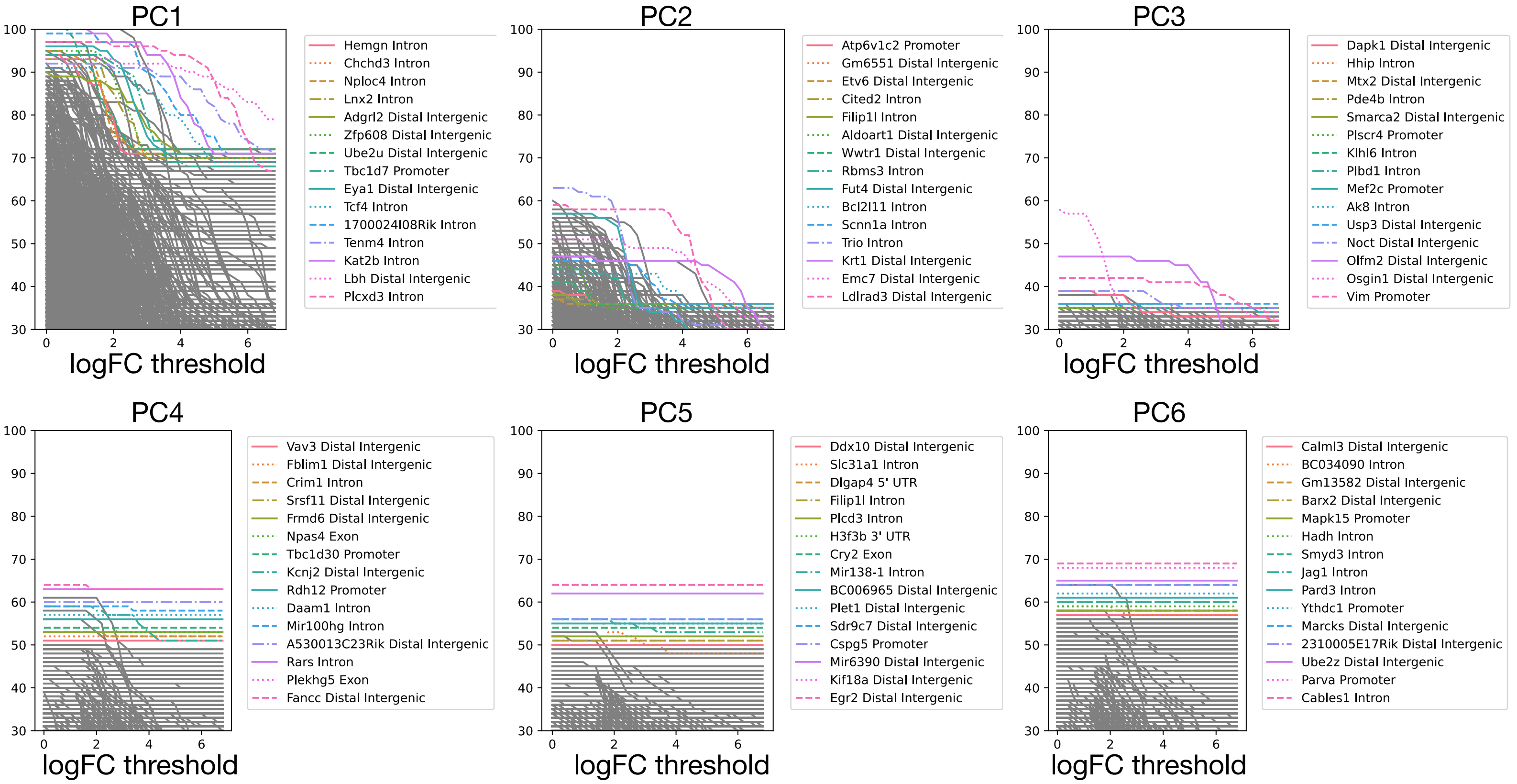


**(b) Top genes explained by principal components of scATAC-seq modality-specific latent space**

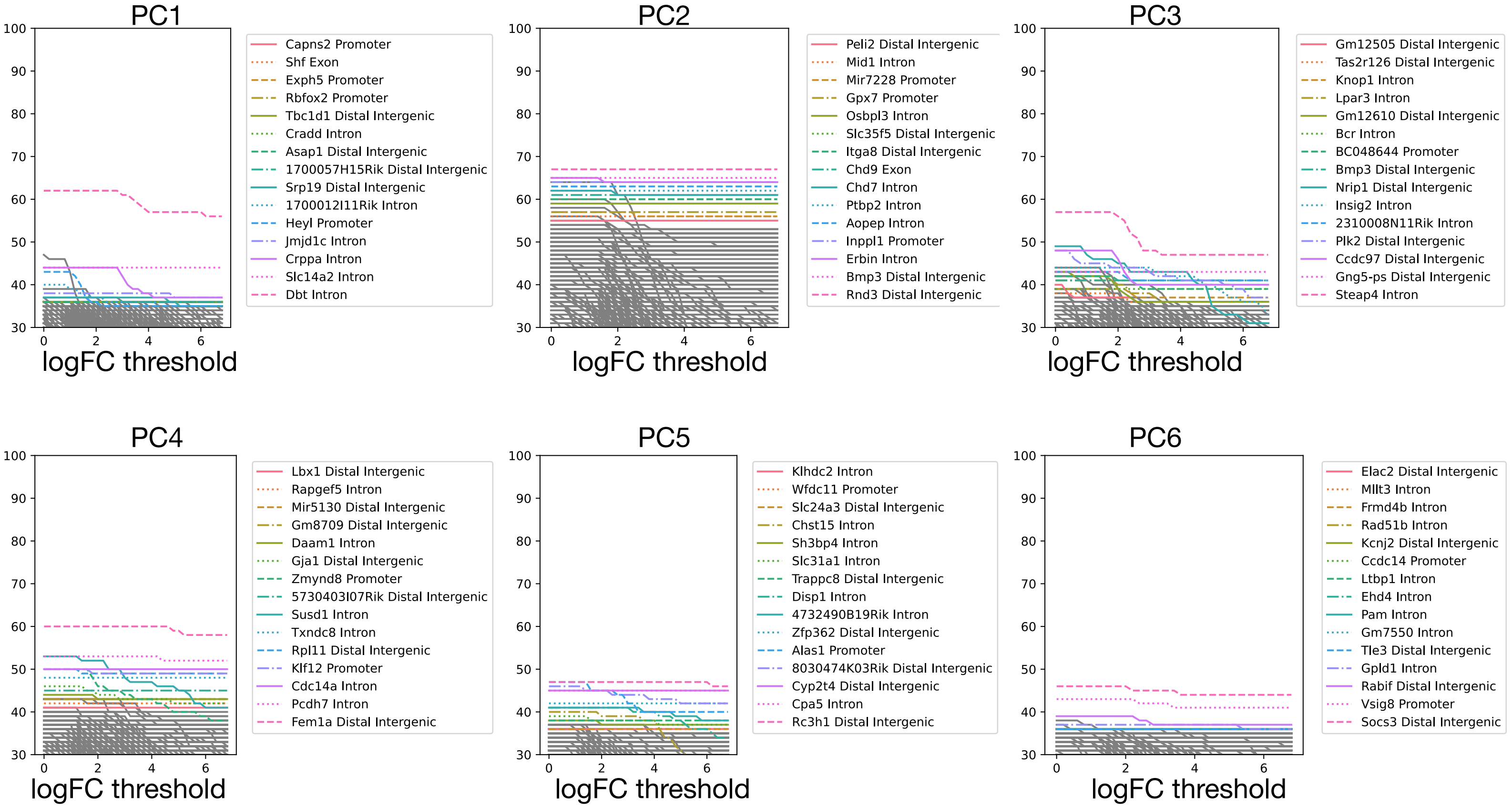


**Figure S4**

**Figure S4. Top genes explained by scATAC-seq shared and modality-specific latent spaces**
We test for significant gene expression and peak count changes along each PC in the (a) scATAC-seq shared latent space and (b) scATAC-seq modality-specific latent space. The cells are divided into 36 random train-test splits with 20% of cells used for testing in each split. The threshold of p-values after multiple testing correction is set to 0.05. The number of times in all train-test splits a gene or peak is tested to be significant in both the training and testing cells with the same direction of fold change is plotted for different thresholds of log fold change magnitude.

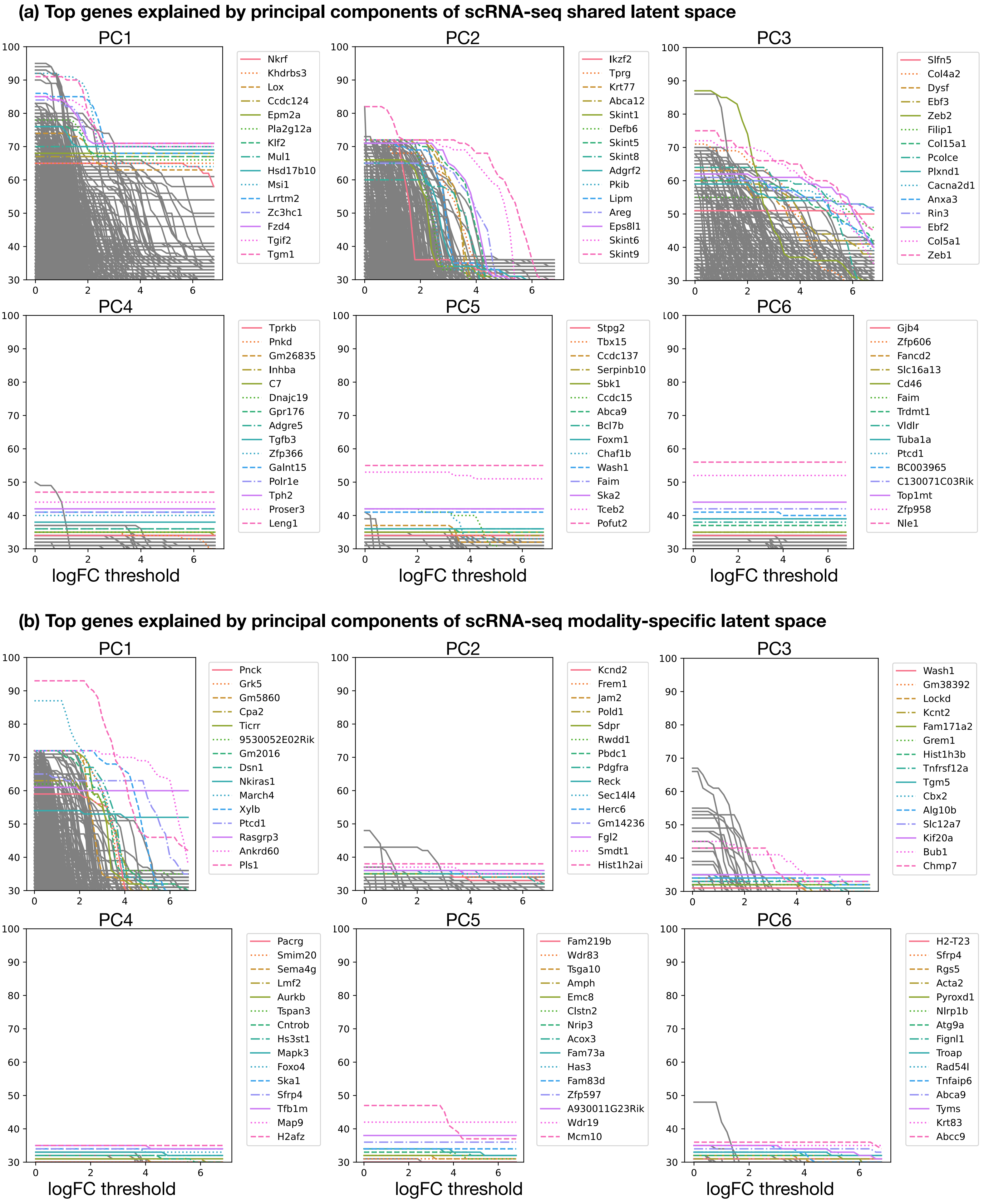
(a) Top genes explained by principal components of scRNA-seq shared latent space
PC1
Nkrf
Khdrbs3
Lox
Ccdc124
Epm2a
Pla2g12a
Klf2
Mul1
Hsd17b10
Msi1
Lrrtm2
Zc3hc1
Fzd4
Tgif2
Tgm1
PC2
Ikzf2
Tprg
Krt77
Abca12
Skint1
Defb6
Skint5
Skint8
Adgrf2
Pkib
Lipm
Areg
Eps8l1
Skint6
Skint9
PC3
Slfn5
Col4a2
Dysf
Ebf3
Zeb2
Filip1
Col15a1
Pcolce
Plxnd1
Cacna2d1
Anxa3
Rin3
Ebf2
Col5a1
Zeb1
PC4
Tprkb
Pnkd
Gm26835
Inhba
C7
Dnajc19
Gpr176
Adgre5
Tgfb3
Zfp366
Galnt15
Polr1e
Tph2
Proser3
Leng1
PC5
Stpg2
Tbx15
Ccdc137
Serpinb10
Sbk1
Ccdc15
Abca9
Bcl7b
Foxm1
Chaf1b
Wash1
Faim
Ska2
Tceb2
Pofut2
PC6
Gjb4
Zfp606
Fancd2
Slc16a13
Cd46
Faim
Trdmt1
Vldlr
Tuba1a
Ptcd1
BC003965
C130071C03Rik
Top1mt
Zfp958
Nle1
logFC threshold
(b) Top genes explained by principal components of scRNA-seq modality-specific latent space
PC1
Pnck
Grk5
Gm5860
Cpa2
Ticrr
9530052E02Rik
Gm2016
Dsn1
Nkiras1
March4
Xylb
Ptcd1
Rasgrp3
Ankrd60
Pls1
PC2
Kcnd2
Frem1
Jam2
Pold1
Sdpr
Rwdd1
Pbdc1
Pdgfra
Reck
Sec14l4
Herc6
Gm14236
Fgl2
Smdt1
Hist1h2ai
PC3
Wash1
Gm38392
Lockd
Kcnt2
Fam171a2
Grem1
Hist1h3b
Tnfrsf12a
Tgm5
Cbx2
Alg10b
Slc12a7
Kif20a
Bub1
Chmp7
PC4
Pacrg
Smim20
Sema4g
Lmf2
Aurkb
Tspan3
Cntrob
Hs3st1
Mapk3
Foxo4
Ska1
Sfrp4
Tfb1m
Map9
H2afz
PC5
Fam219b
Wdr83
Tsga10
Amph
Emc8
Clstn2
Nrip3
Acox3
Fam73a
Has3
Fam83d
Zfp597
A930011G23Rik
Wdr19
Mcm10
PC6
H2-T23
Sfrp4
Rgs5
Acta2
Pyroxd1
Nlrp1b
Atg9a
Fignl1
Troap
Rad54l
Tnfaip6
Abca9
Tyms
Krt83
Abcc9
logFC threshold

**Figure S5**

**Figure S5. Top genes explained by scRNA-seq shared and modality-specific latent spaces**
We test for significant gene expression and peak count changes along each PC in the (a) scRNA-seq shared latent space and (b) scRNA-seq modality-specific latent space. The cells are divided into 36 random train-test splits with 20% of cells used for testing in each split. The threshold of p-values after multiple testing correction is set to 0.05. The number of times in all train-test splits a gene or peak is tested to be significant in both the training and testing cells with the same direction of fold change is plotted for different thresholds of log fold change magnitude.

| Patient ID | Diagnosis | Train-test | DE_heldout_group |
|---|---|---|---|
| HV1 | healthy | Test | 1, 5, 9 |
| HV2 | healthy | Train | 3, 5, 10 |
| HV3 | healthy | Train | 1, 6, 10 |
| HV4 | healthy | Train | 4, 6, 11 |
| HV5 | healthy | Train | 2, 7, 9 |
| HV6 | healthy | Train | 4, 7, 12 |
| HV7 | healthy | Train | 2, 8, 11 |
| HV8 | healthy | Train | 3, 8, 12 |
| HV9 | healthy | Train | |
| HV10 | healthy | Train | |
| P27 | Meningioma | Test | 1, 5, 10 |
| P33 | Meningioma | Train | |
| P37 | Meningioma | Train | 1, 6, 11 |
| P38 | Meningioma | Train | 4, 9 |
| P42 | Meningioma | Train | 2, 6, 12 |
| P48 | Meningioma | Train | 5, 9 |
| P59 | Meningioma | Train | 2, 7, 10 |
| P62 | Meningioma | Train | 4, 8 |
| P70 | Meningioma | Train | 3, 7, 11 |
| P83 | Meningioma | Train | 3, 8, 12 |
| P15 | Glioma | Train | 2, 7, 12 |
| P16 | Glioma | Train | 4, 8, 11 |
| P22 | Glioma | Test | 1, 5, 10 |
| P29 | Glioma | Train | |
| P46 | Glioma | Train | 1, 6, 12 |
| P47 | Glioma | Train | 3, 7, 11 |
| P52 | Glioma | Train | 4, 9 |
| P57 | Glioma | Train | 3, 8 |
| P68 | Glioma | Train | 2, 6, 10 |
| P84 | Glioma | Train | 5, 9 |
| P14 | Headneck | Test | 1, 6, 12 |
| P18 | Headneck | Train | 4, 8 |
| P24 | Headneck | Train | 1, 7, 10 |
| P41 | Headneck | Train | 4, 9 |
| P44 | Headneck | Train | 2, 6, 11 |
| P50 | Headneck | Train | 5, 10 |
| P55 | Headneck | Train | 2, 7, 12 |
| P56 | Headneck | Train | 5, 11 |
| P63 | Headneck | Train | 3, 8 |
| P72 | Headneck | Train | 3, 9 |

**Figure S6**

**Figure S6. Train-test split of the PBMC dataset with paired chromatin and protein images.**
First two columns list the patient IDs and the clinical diagnosis. "Train-test" column lists whether the cells in a particular patient are used to train the APOLLO model or held-out for testing. "DE_heldout_group" lists the train-test splits used for identifying significant morphological feature changes along each PC of the image latent spaces. There are a total of 12 different train-test splits and the numbers indicate in which splits the particular patient is in the test group.

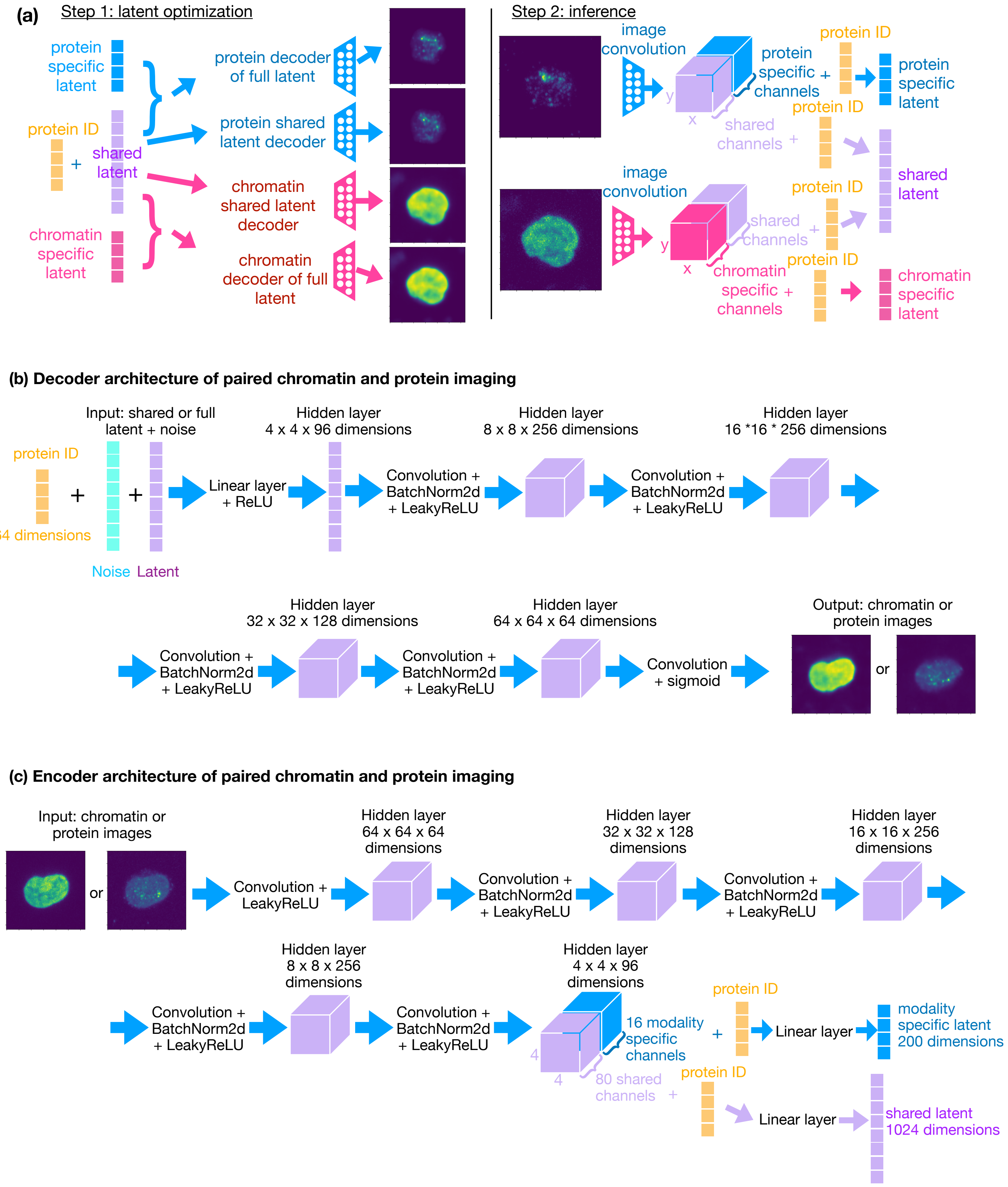

(a)
Step 1: latent optimization
protein specific latent
protein ID
+
shared latent
chromatin specific latent
protein decoder of full latent
protein shared latent decoder
chromatin shared latent decoder
chromatin decoder of full latent
Step 2: inference
image convolution
protein specific + channels
shared channels +
protein ID
protein specific latent
shared latent
image convolution
shared channels +
chromatin specific + channels
chromatin specific latent
x
y
(b) Decoder architecture of paired chromatin and protein imaging
Input: shared or full latent + noise
protein ID
64 dimensions
Noise
Latent
Linear layer + ReLU
Hidden layer 4 x 4 x 96 dimensions
Convolution + BatchNorm2d + LeakyReLU
Hidden layer 8 x 8 x 256 dimensions
Hidden layer 16 *16 * 256 dimensions
Hidden layer 32 x 32 x 128 dimensions
Hidden layer 64 x 64 x 64 dimensions
Convolution + sigmoid
Output: chromatin or protein images
or
(c) Encoder architecture of paired chromatin and protein imaging
Input: chromatin or protein images
Convolution + LeakyReLU
Hidden layer 64 x 64 x 64 dimensions
Convolution + BatchNorm2d + LeakyReLU
Hidden layer 32 x 32 x 128 dimensions
Hidden layer 16 x 16 x 256 dimensions
Hidden layer 8 x 8 x 256 dimensions
Hidden layer 4 x 4 x 96 dimensions
16 modality specific channels
80 shared channels
protein ID
Linear layer
modality specific latent 200 dimensions
shared latent 1024 dimensions


**Figure S7**

**Figure S7. APOLLO architecture for paired chromatin and protein images.** (a) APOLLO can be applied to multiplexed imaging data using conditional decoders and encoders. In step 1, the shared and modality-specific latent spaces as well as the modality-specific decoders are trained to minimize reconstruction error of chromatin and protein images from the latent spaces. In addition to the latent space embeddings, the decoders also take in a trainable vector representing each protein ID. In step 2, modality-specific encoders are trained to enable inferring the latent space embedding for cells not used in training the model and thus enable cross modality prediction. The encoders are trained to map protein and chromatin images together with the trainable protein IDs to the latent space embeddings obtained in step 1.
(b) The decoders of chromatin and protein images have the same architectures with five convolutional layers following a fully connected layer. The input to the decoder is the full or shared latent space added with noise and concatenated with learnable protein ID. The latent space embedding, after adding noise and being concatenated with the protein ID, is passed through a linear layer with ReLU activation and reshaped to $4 \times 4 \times 96$ dimensions for subsequent convolutions. This is followed by 5 convolutional layers with a kernel size of 4 and stride of 2. The first four convolutional layers are followed by batch normalization and LeakyReLU activation. The last convolutional layer is followed by sigmoid activation to scale the output image from 0 to 1.
(c) The encoder of each modality starts with 5 convolutional layers with LeakyReLU activation, a standard setup for autoencoders [3, 9, 15]. The output of the last convolutional layer is divided into two sets of channels that are used to derive the shared and modality-specific latent spaces respectively. 80 out of the 96 channels of the last hidden layer are flattened, concatenated with protein ID, and passed through a linear layer to obtain the shared latent space. Similarly, the modality-specific latent space is obtained from the remaining 16 channels.

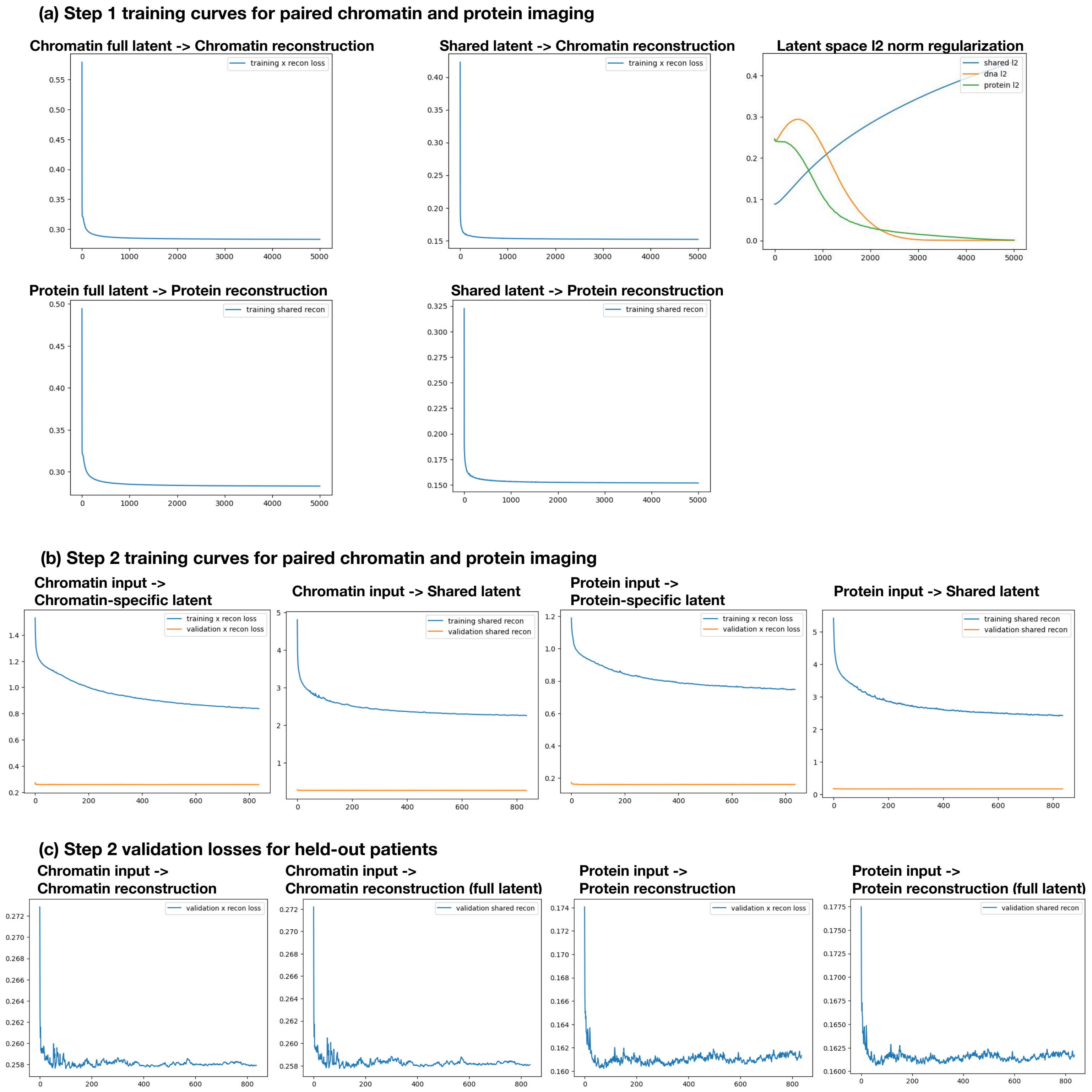
(a) Step 1 training curves for paired chromatin and protein imaging
Chromatin full latent -> Chromatin reconstruction
training x recon loss
Shared latent -> Chromatin reconstruction
training x recon loss
Latent space l2 norm regularization
shared l2
dna l2
protein l2
Protein full latent -> Protein reconstruction
training shared recon
Shared latent -> Protein reconstruction
training shared recon
(b) Step 2 training curves for paired chromatin and protein imaging
Chromatin input -> Chromatin-specific latent
training x recon loss
validation x recon loss
Chromatin input -> Shared latent
training shared recon
validation shared recon
Protein input -> Protein-specific latent
training x recon loss
validation x recon loss
Protein input -> Shared latent
training shared recon
validation shared recon
(c) Step 2 validation losses for held-out patients
Chromatin input -> Chromatin reconstruction
validation x recon loss
Chromatin input -> Chromatin reconstruction (full latent)
validation shared recon
Protein input -> Protein reconstruction
validation x recon loss
Protein input -> Protein reconstruction (full latent)
validation shared recon

**Figure S8**

**Figure S8. Training and validation curves of APOLLO applied to paired chromatin and protein images.**

(a) Training curves are plotted for the latent optimization in step 1. "Chromatin full latent → Chromatin reconstruction" shows the chromatin reconstruction loss using the full chromatin latent space, i.e., the shared latent space embedding concatenated with the chromatin-specific latent space embedding. "Shared latent → Chromatin reconstruction" whose the chromatin reconstruction loss using the shared latent space. "Protein full latent → Protein reconstruction" shows the protein reconstruction loss using the full protein latent space, i.e., the shared latent space embedding concatenated with the protein-specific latent space embedding. "Shared latent → Protein reconstruction" shows the protein reconstruction loss using the shared latent space. "Latent space l2 norm regularization" shows the $\ell_2$ norms of the three latent spaces during training.

(b) Training curves for the inference in step 2 are plotted for the reconstruction of the shared and modality-specific latent spaces learned in step 1. "Chromatin input → Chromatin-specific latent" considers the reconstruction of the chromatin-specific latent space from chromatin images using the chromatin-encoder. "Chromatin input → Shared latent" considers the reconstruction of the shared latent space from chromatin images using the chromatin-encoder. "Protein input → Protein-specific latent" considers the reconstruction of the protein-specific latent space from protein images using the protein-encoder. "Protein input → Shared latent" considers the reconstruction of the shared latent space from protein images using the protein-encoder.

(c) Chromatin and protein reconstruction losses are plotted for cells from patients held out during training. "Chromatin input → chromatin reconstruction" shows the reconstruction loss of held-out cells obtained from first inferring the shared latent space using the chromatin encoder and then decoding the chromatin images using the chromatin decoder of the shared latent space. "Chromatin input → Chromatin reconstruction (full latent)" shows the reconstruction loss of held-out cells obtained from first inferring the shared latent space and the chromatin-specific latent space using the chromatin encoder and then decoding the chromatin images using the chromatin decoder of the full chromatin latent space. "Protein input → Protein reconstruction" shows the reconstruction loss of held-out cells obtained from first inferring the shared latent space using the protein encoder and then decoding the protein images using the protein decoder of the shared latent space. "Protein input → protein reconstruction (full latent)" shows the reconstruction loss of held-out cells obtained from first inferring the shared latent space and the protein-specific latent space using the protein encoder and then decoding the protein images using the protein decoder of the full protein latent space.

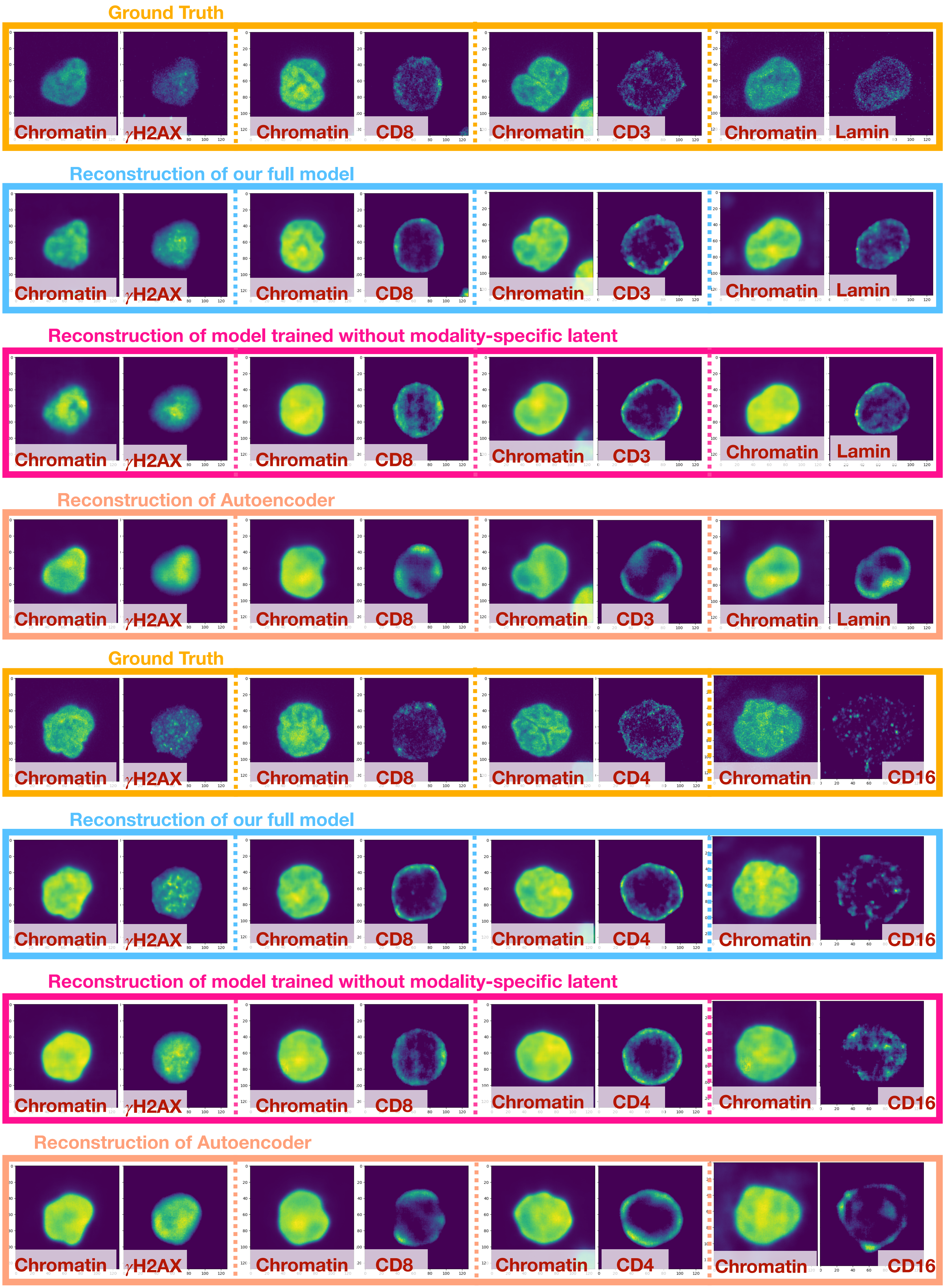

Ground Truth
Chromatin
γH2AX
Chromatin
CD8
Chromatin
CD3
Chromatin
Lamin
Reconstruction of our full model
Chromatin
γH2AX
Chromatin
CD8
Chromatin
CD3
Chromatin
Lamin
Reconstruction of model trained without modality-specific latent
Chromatin
γH2AX
Chromatin
CD8
Chromatin
CD3
Chromatin
Lamin
Reconstruction of Autoencoder
Chromatin
γH2AX
Chromatin
CD8
Chromatin
CD3
Chromatin
Lamin
Ground Truth
Chromatin
γH2AX
Chromatin
CD8
Chromatin
CD4
Chromatin
CD16
Reconstruction of our full model
Chromatin
γH2AX
Chromatin
CD8
Chromatin
CD4
Chromatin
CD16
Reconstruction of model trained without modality-specific latent
Chromatin
γH2AX
Chromatin
CD8
Chromatin
CD4
Chromatin
CD16
Reconstruction of Autoencoder
Chromatin
γH2AX
Chromatin
CD8
Chromatin
CD4
Chromatin
CD16


**Figure S9**

**Figure S9. Examples of reconstructed chromatin and protein images of cells from held-out patients obtained from our full APOLLO model are compared to the outputs of different alternative models.**
"Autoencoder" has the same encoder and decoder structures as our full APOLLO model but is trained in a single step without directly optimizing the latent spaces (Extended Methods). "Model trained without modality-specific latent" represents our APOLLO model with only the shared latent space.

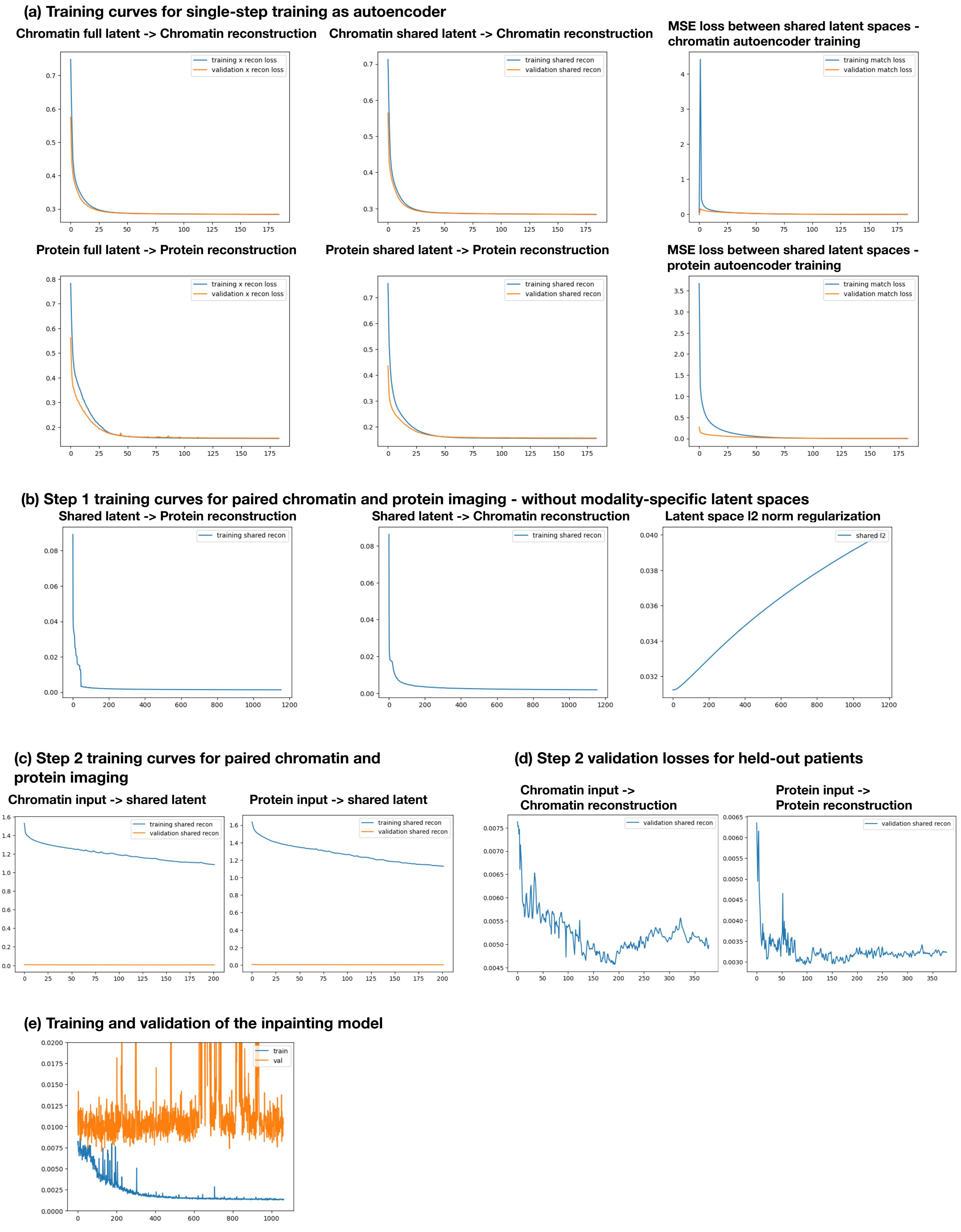
(a) Training curves for single-step training as autoencoder
Chromatin full latent -> Chromatin reconstruction
training x recon loss
validation x recon loss
Chromatin shared latent -> Chromatin reconstruction
training shared recon
validation shared recon
MSE loss between shared latent spaces - chromatin autoencoder training
training match loss
validation match loss
Protein full latent -> Protein reconstruction
training x recon loss
validation x recon loss
Protein shared latent -> Protein reconstruction
training shared recon
validation shared recon
MSE loss between shared latent spaces - protein autoencoder training
training match loss
validation match loss
(b) Step 1 training curves for paired chromatin and protein imaging - without modality-specific latent spaces
Shared latent -> Protein reconstruction
training shared recon
Shared latent -> Chromatin reconstruction
training shared recon
Latent space l2 norm regularization
shared l2
(c) Step 2 training curves for paired chromatin and protein imaging
Chromatin input -> shared latent
training shared recon
validation shared recon
Protein input -> shared latent
training shared recon
validation shared recon
(d) Step 2 validation losses for held-out patients
Chromatin input -> Chromatin reconstruction
validation shared recon
Protein input -> Protein reconstruction
validation shared recon
(e) Training and validation of the inpainting model
train
val

Figure S10

**Figure S10. Training and validation curves of models used for benchmarking.** (a) The plots show training and validation curves of the model that has the same encoder and decoder structures as our full APOLLO model but is trained in a single step without directly optimizing the latent spaces (Extended Methods). "Chromatin full latent → Chromatin reconstruction" shows the reconstruction loss computed using the chromatin decoder of the full chromatin latent space. "Chromatin shared latent → Chromatin reconstruction" shows the reconstruction loss computed using the chromatin decoder of the shared latent space encoded by the chromatin encoder. "MSE loss between shared latent spaces - chromatin autoencoder training" and "MSE loss between shared latent spaces - protein autoencoder training" show the MSE loss used to match the shared latent spaces encoded by the chromatin encoder and by the protein encoder during training. Since we alternate between training of the chromatin and the protein autoencoders in each epoch, the two plots show the MSE loss computed during each autoencoder training separately. "Protein full latent → Protein reconstruction" shows the reconstruction loss computed using the protein decoder of the full protein latent space. "Protein shared latent → Protein reconstruction" shows the reconstruction loss computed using the protein decoder of the shared latent space encoded by the protein encoder.
(b) Training curves are plotted for the latent optimization in step 1 of our model without modality-specific latent spaces. "Shared latent → Chromatin reconstruction" shows the chromatin reconstruction loss using the shared latent space. "Shared latent → Protein reconstruction" shows the protein reconstruction loss using the shared latent space. "Latent space l2 norm regularization" shows the $\ell_2$ norms of the three latent spaces during training.
(c) Training curves for the inference in step 2 are plotted for the reconstruction of the shared and modality-specific latent spaces learned in step 1. "Chromatin input → Shared latent" considers the reconstruction of the shared latent space from chromatin images using the chromatin-encoder. "Protein input → Shared latent" considers the reconstruction of the shared latent space from protein images using the protein-encoder.
(d) Chromatin and protein reconstruction losses are plotted for cells from patients held out during training. "Chromatin input → chromatin reconstruction" shows the reconstruction loss of held-out cells obtained from first inferring the shared latent space using the chromatin encoder and then decoding the chromatin images using the chromatin decoder of the shared latent space. "Protein input → Protein reconstruction" shows the reconstruction loss of held-out cells obtained from first inferring the shared latent space using the protein encoder and then decoding the protein images using the protein decoder of the shared latent space.
(e) Training and validation curves obtained from training the inpainting model from [9].

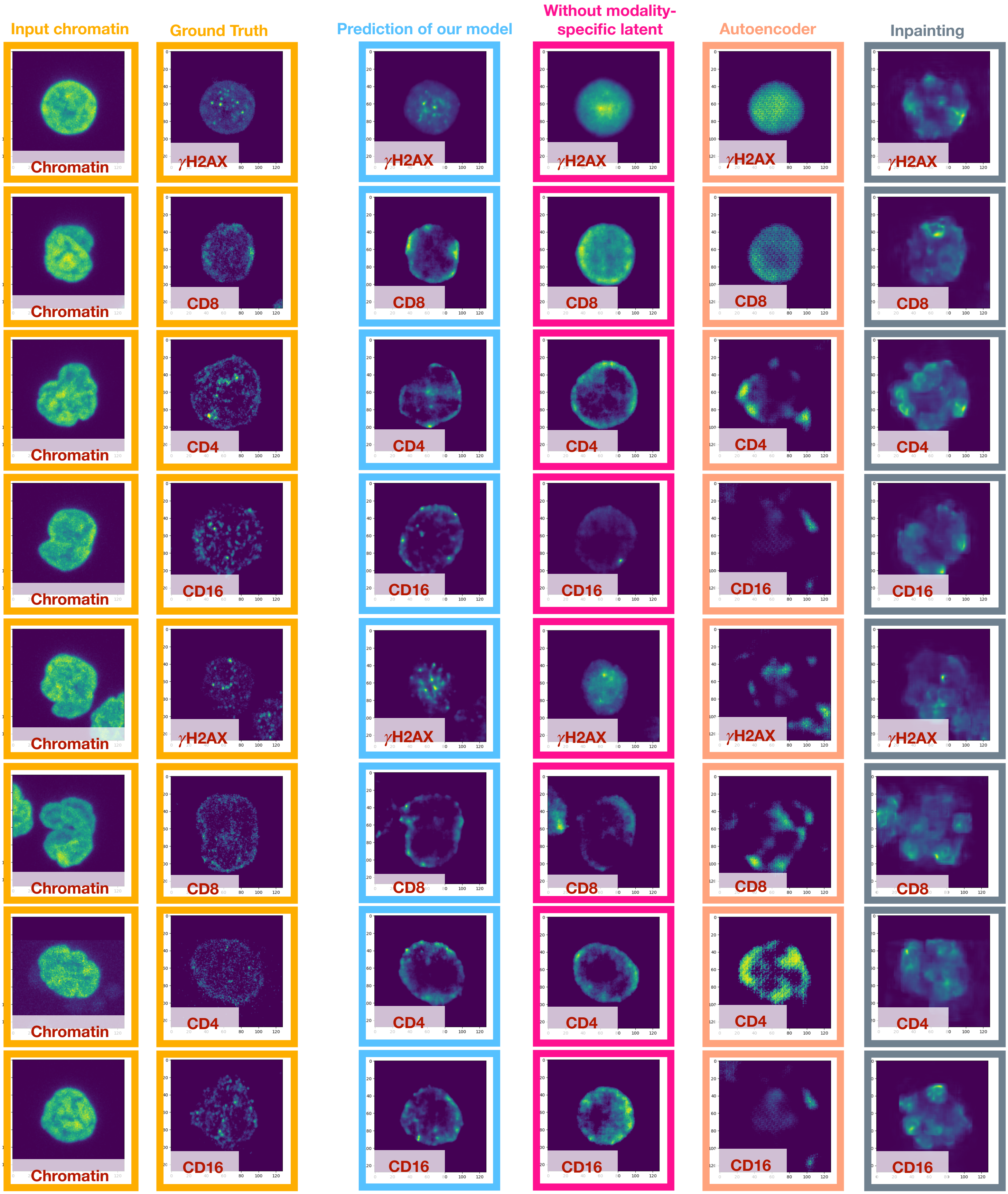
Input chromatin
Ground Truth
Prediction of our model
Without modality-specific latent
Autoencoder
Inpainting
Chromatin
γH2AX
CD8
CD4
CD16

**Figure S11**

**Figure S11. Examples of protein images predicted from chromatin images of held-out patients obtained from our full APOLLO model are compared to the outputs of different alternative models.**
"Autoencoder" has the same encoder and decoder structures as our full APOLLO model but is trained in a single step without directly optimizing the latent spaces (Extended Methods). "Without modality-specific latent" represents our APOLLO model with only the shared latent space. "Inpainting" corresponds to the model developed by [9].

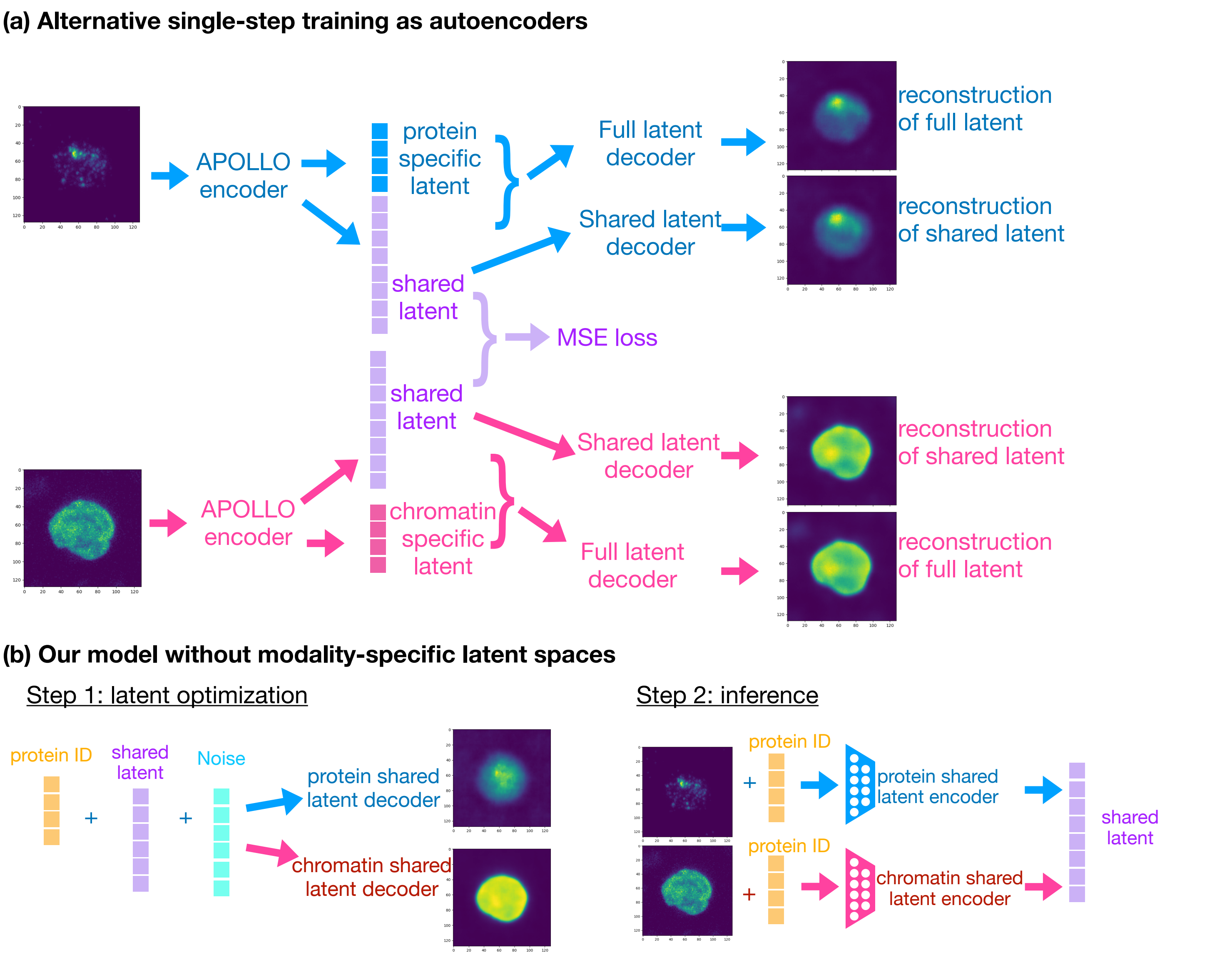
(a) Alternative single-step training as autoencoders
APOLLO encoder
protein specific latent
Full latent decoder
reconstruction of full latent
Shared latent decoder
reconstruction of shared latent
shared latent
MSE loss
shared latent
Shared latent decoder
reconstruction of shared latent
APOLLO encoder
chromatin specific latent
Full latent decoder
reconstruction of full latent
(b) Our model without modality-specific latent spaces
Step 1: latent optimization
protein ID
shared latent
Noise
protein shared latent decoder
chromatin shared latent decoder
Step 2: inference
protein ID
protein shared latent encoder
protein ID
chromatin shared latent encoder
shared latent

**Figure S12**

**Figure S12. Architecture details of model ablation tests and comparison of reconstruction loss.**
(a) The schematic shows the architecture of our model with partially overlapping latent space trained as a standard autoencoder in a single step. The encoders and decoders have the same architecture as our APOLLO model, but the encoder and decoder are trained together in a single step without directly optimizing the latent spaces. Mean-squared error loss is applied to the shared latent space encoded from each modality.
(b) The schematic shows the architecture of our model without the modality-specific latent spaces. The model is trained with the two-step training procedure as used for the full APOLLO model. There are only two decoders that decode the two modalities from the shared latent space and the are trained in step 1. The encoders only output the shared latent space, which has the same dimension as the combined dimension of the shared and the modality-specific latent spaces in the full APOLLO model.

**(a) Representative features with significant changes along the PCs - Chromatin shared latent space**

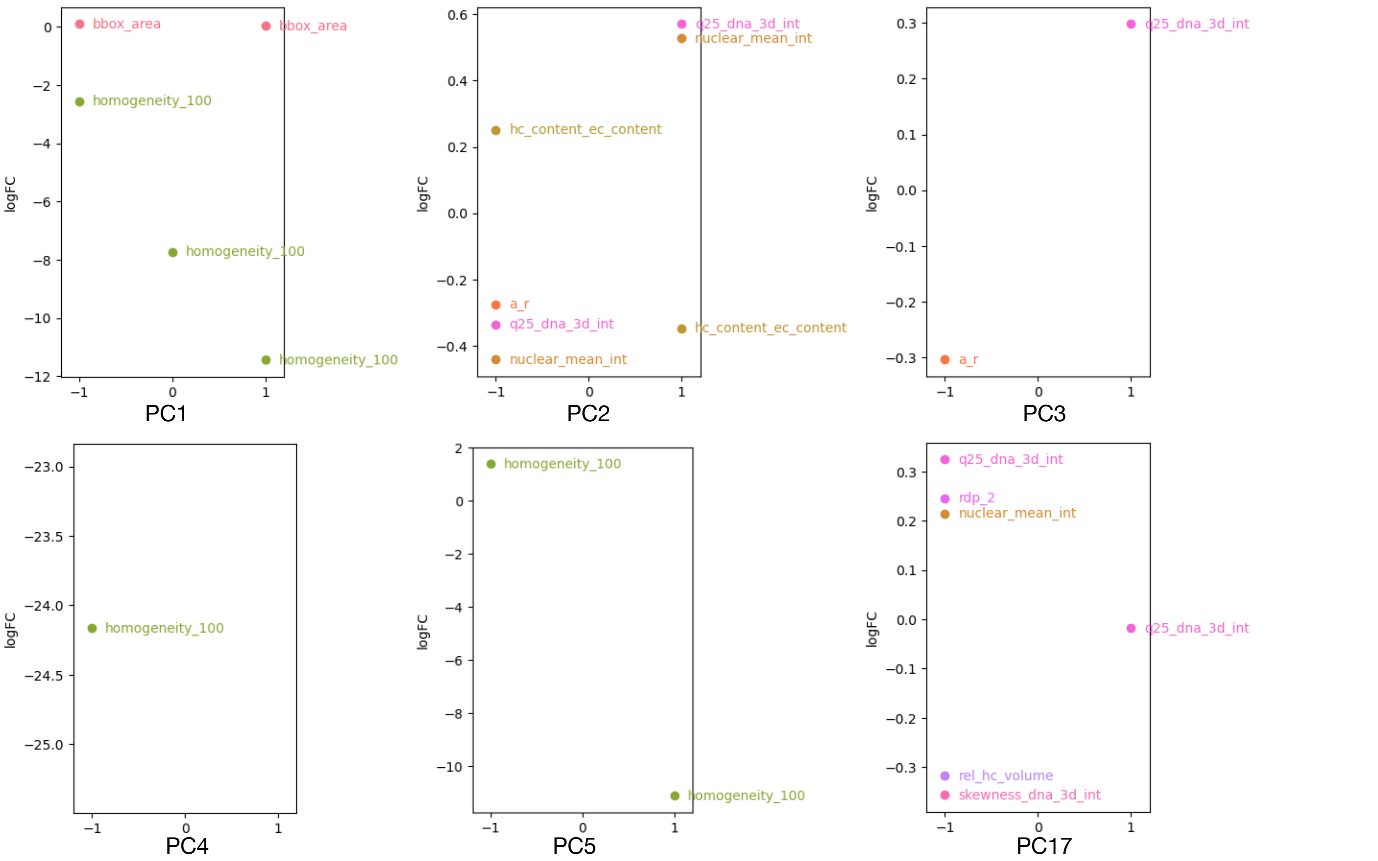


**(b) Representative features with significant changes along the PCs - Chromatin modality-specific latent space**

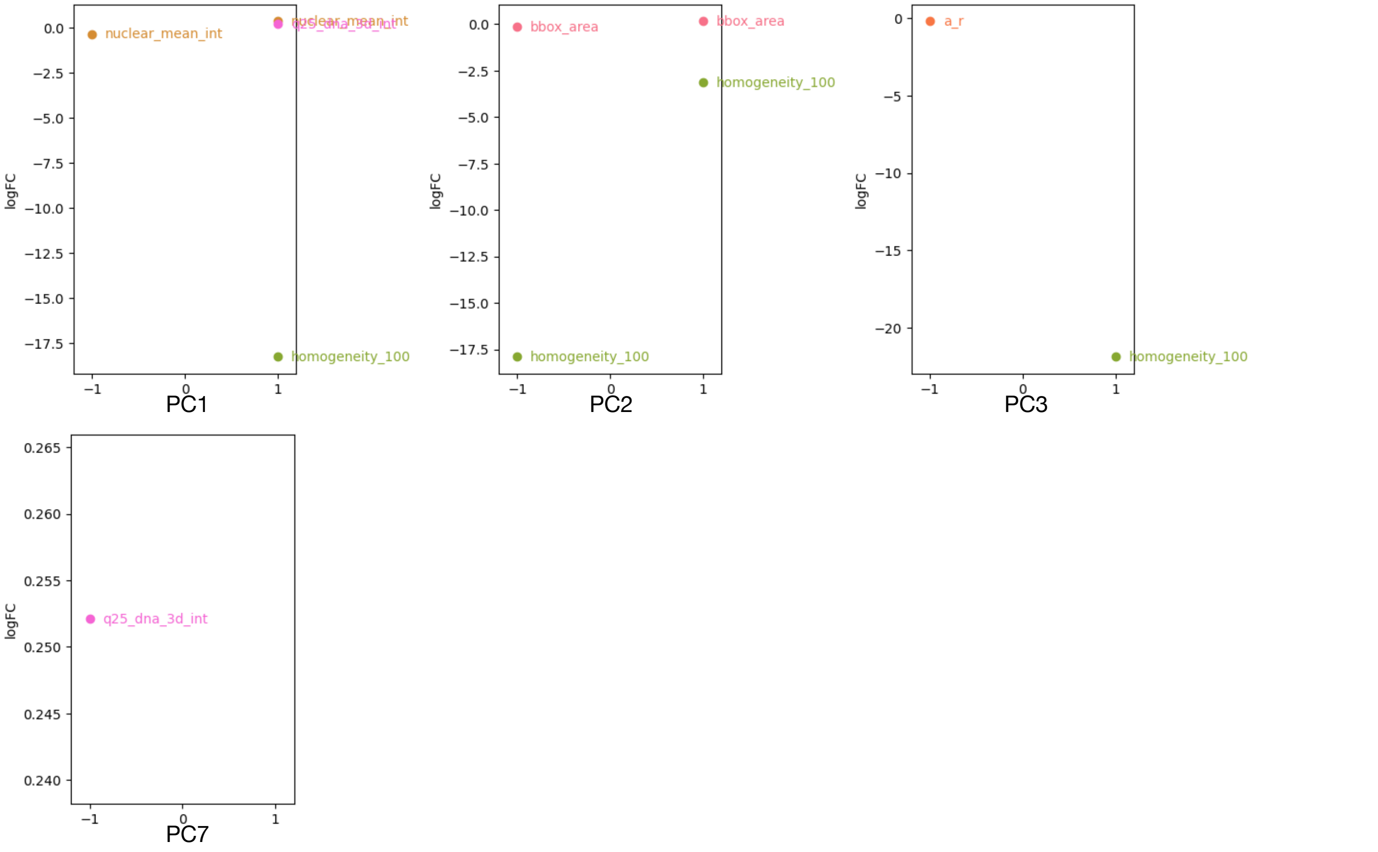


**Figure S13**

**Figure S13. Representative features in the chromatin shared and modality-specific latent spaces.** Only one feature from each group of highly correlated features is plotted. The approach to feature grouping and selection of representative features is described in Extended Methods. As described in the Extended Methods section, we consider three groups of cells for each $PC_i$ that are all at the center of other PCs, which are labeled as -1, 0, and 1 respectively.

**(a) Representative features with significant changes along the PCs - CD16 shared latent space**

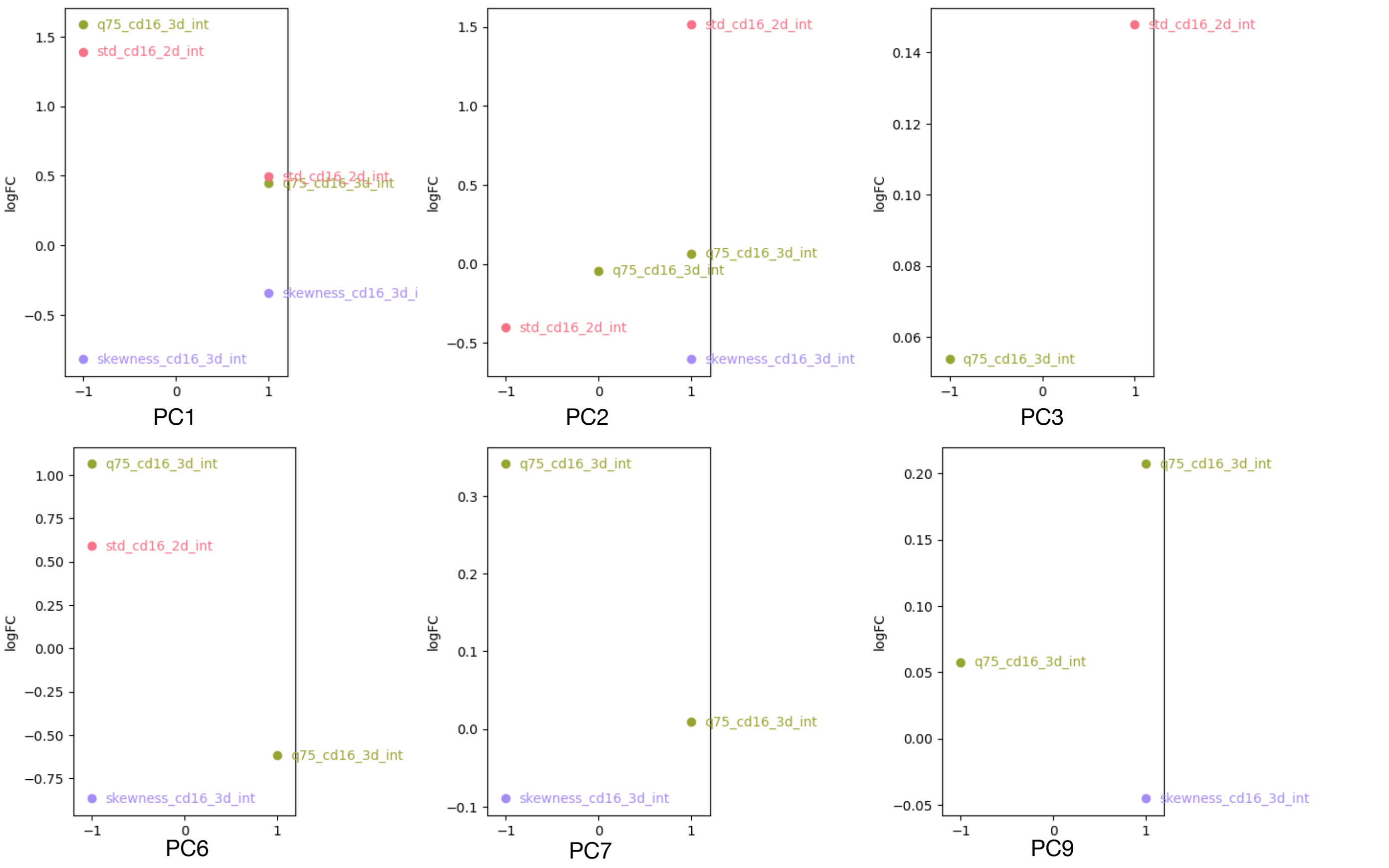


**(b) Representative features with significant changes along the PCs - CD16 modality-specific latent space**

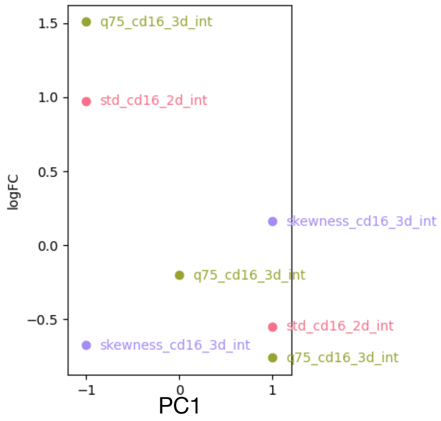


**(c) Summary of the morphological features explained by each latent space**

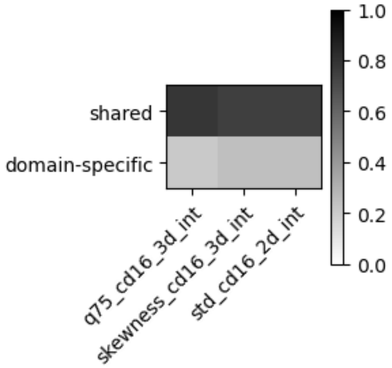


**Figure S14**

**Figure S14. Representative features in the CD16 shared and modality-specific latent spaces.** Only one feature from each group of highly correlated features is plotted. The approach to feature grouping and selection of representative features is described in Extended Methods. As described in the Extended Methods section, we consider three groups of cells for each $PC_i$ that are all at the center of other PCs, which are labeled as -1, 0, and 1 respectively.

**(a) Representative features with significant changes along the PCs - CD3 shared latent space**

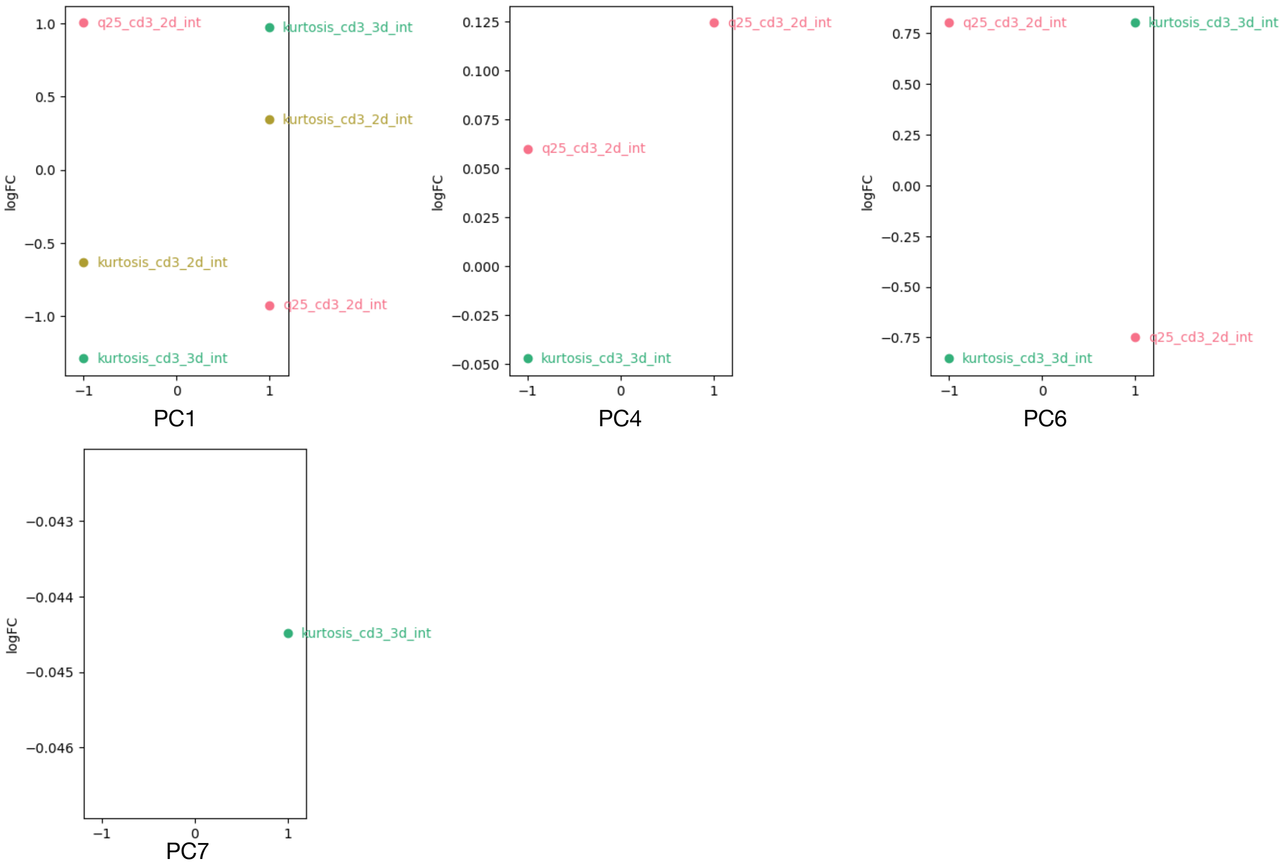


**(b) Representative features with significant changes along the PCs - CD3 modality-specific latent space**

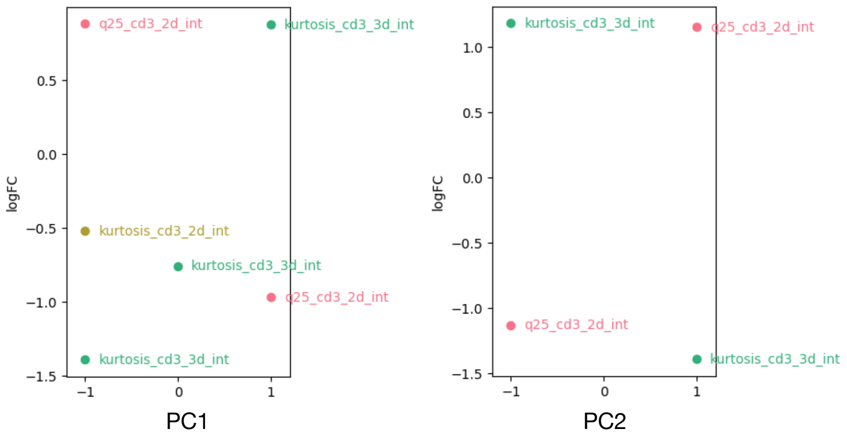


**(c) Summary of the morphological features explained by each latent space**

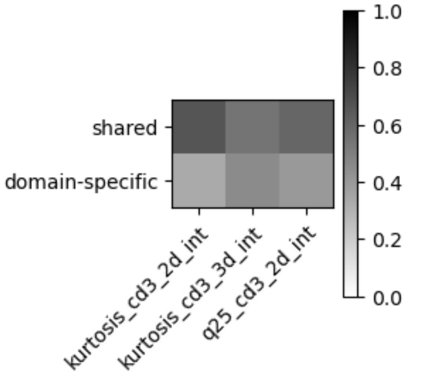


**Figure S15**

**Figure S15. Representative features in the CD3 shared and modality-specific latent spaces.** Only one feature from each group of highly correlated features is plotted. The approach to feature grouping and selection of representative features is described in Extended Methods section. As described in the Extended Methods section, we consider three groups of cells for each $PC_i$ that are all at the center of other PCs, which are labeled as -1, 0, and 1 respectively.

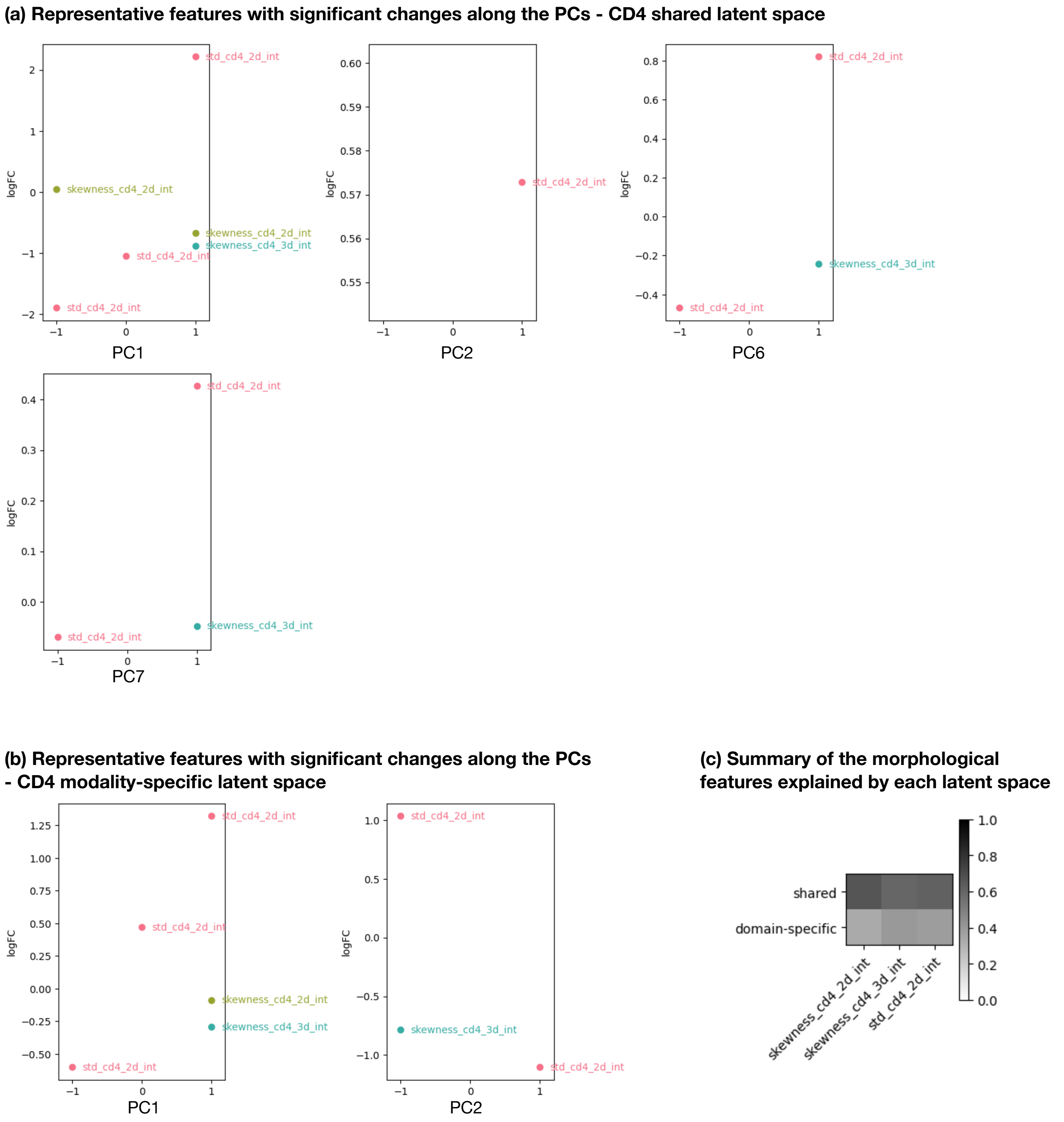
(a) Representative features with significant changes along the PCs - CD4 shared latent space
logFC
PC1
PC2
PC6
PC7
std_cd4_2d_int
skewness_cd4_2d_int
skewness_cd4_3d_int
(b) Representative features with significant changes along the PCs - CD4 modality-specific latent space
(c) Summary of the morphological features explained by each latent space
shared
domain-specific
1.0
0.8
0.6
0.4
0.2
0.0

Figure S16

**Figure S16. Representative features in the CD4 shared and modality-specific latent spaces.** Only one feature from each group of highly correlated features is plotted. The approach to feature grouping and selection of representative features is described in Extended Methods section. As described in the Extended Methods section, we consider three groups of cells for each $PC_i$ that are all at the center of other PCs, which are labeled as -1, 0, and 1 respectively.

**(a) Representative features with significant changes along the PCs - CD8 shared latent space**

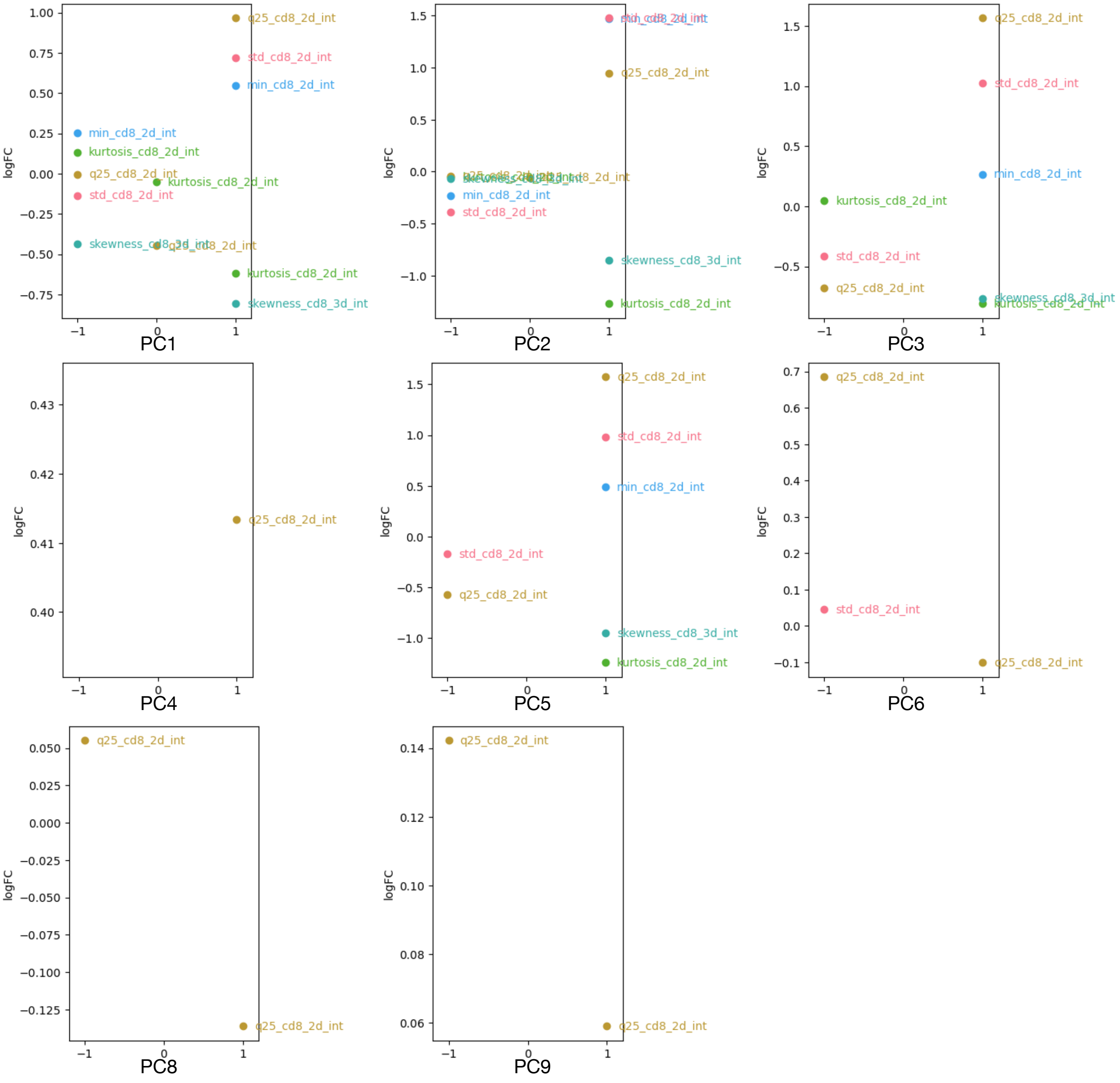


**(b) Representative features with significant changes along the PCs - CD8 modality-specific latent space**

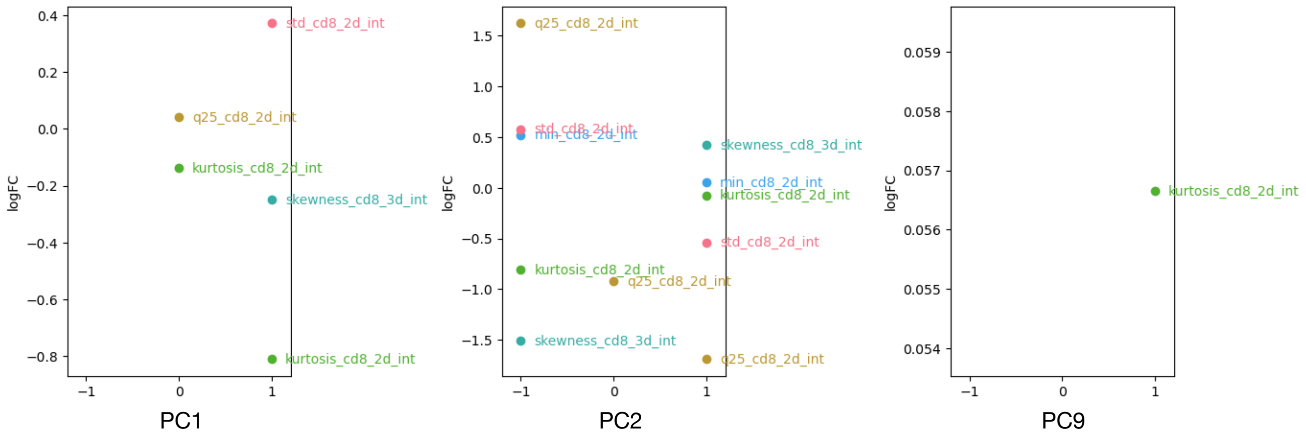


**Figure S17**

**Figure S17. Representative features in the CD8 shared and modality-specific latent spaces.** Only one feature from each group of highly correlated features is plotted. The approach to feature grouping and selection of representative features is described in Extended Methods section. As described in the Extended Methods section, we consider three groups of cells for each $PC_i$ that are all at the center of other PCs, which are labeled as -1, 0, and 1 respectively.

**(a) Representative features with significant changes along the PCs - $\gamma$H2AX shared latent space**

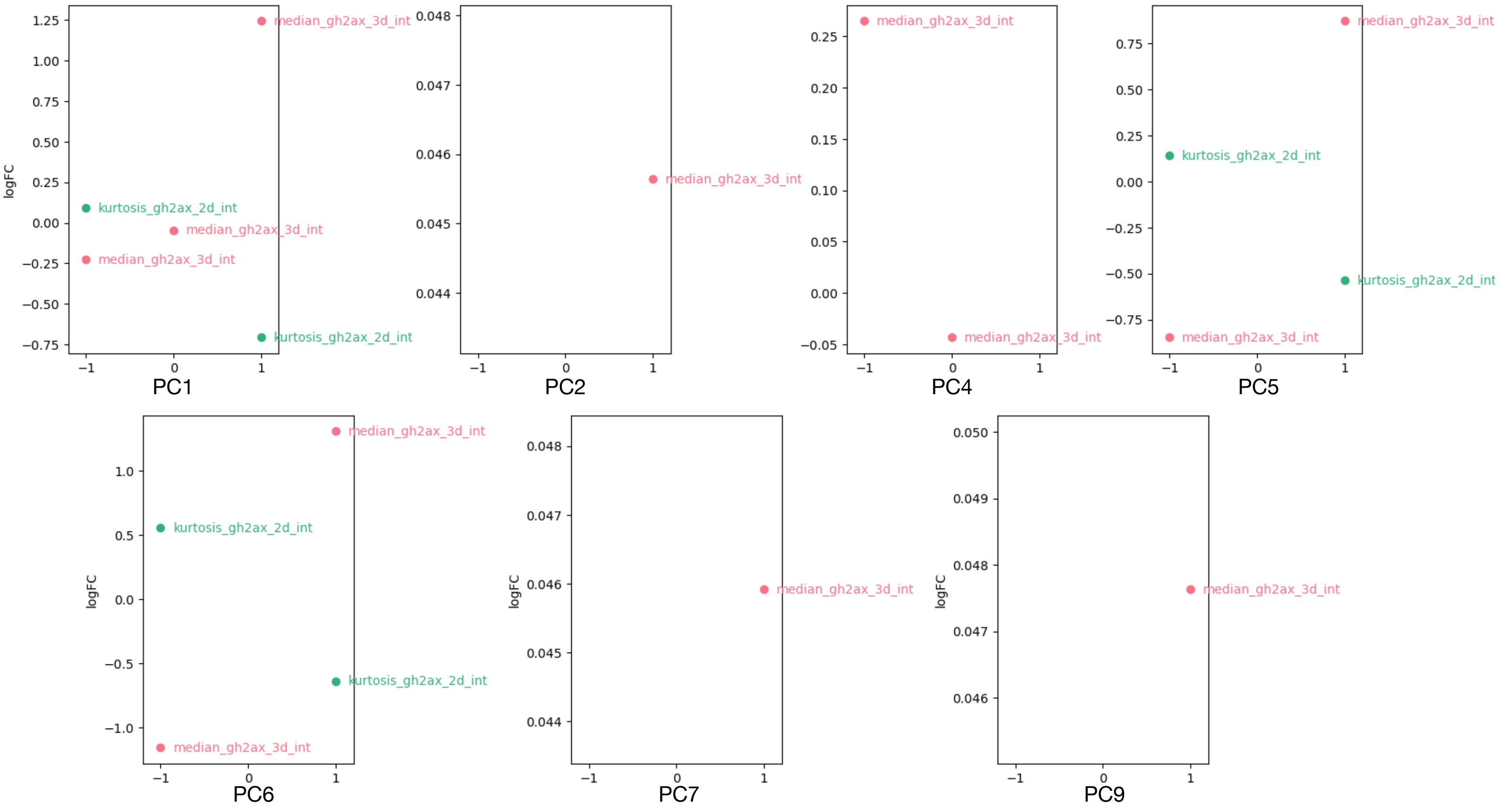


**(b) Representative features with significant changes along the PCs - $\gamma$H2AX modality-specific latent space**

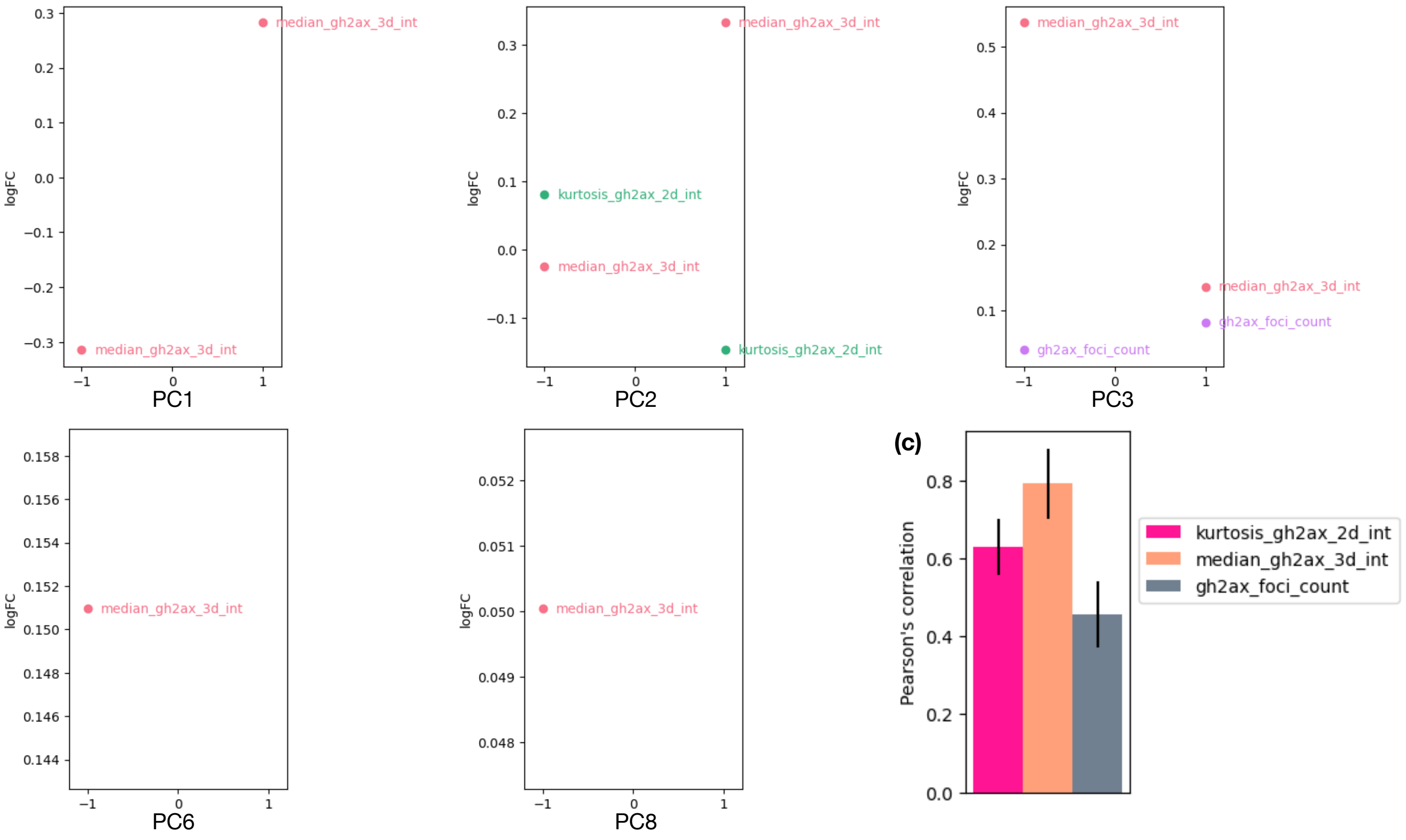


**Figure S18**

**Figure S18. Representative features in the $\gamma$H2AX shared and modality-specific latent spaces.**
(a)-(b) Only one feature from each group of highly correlated features is plotted. The approach to feature grouping and selection of representative features is described in Extended Methods section. As described in the Extended Methods section, we consider three groups of cells for each $PC_i$ that are all at the center of other PCs, which are labeled as -1, 0, and 1 respectively.
(c) Pearson's correlation between the real protein feature values and the predicted values using paired chromatin images. $\gamma$H2AX_foci_count is the only feature that is only captured by the modality-specific space of $\gamma$H2AX but not by chromatin imaging and has the lowest prediction performance ($p = 4.12e-5$ compared to kurtosis_$\gamma$H2AX_2d_int and $p = 8.05e-9$ compared to median_$\gamma$H2AX_3d_int).

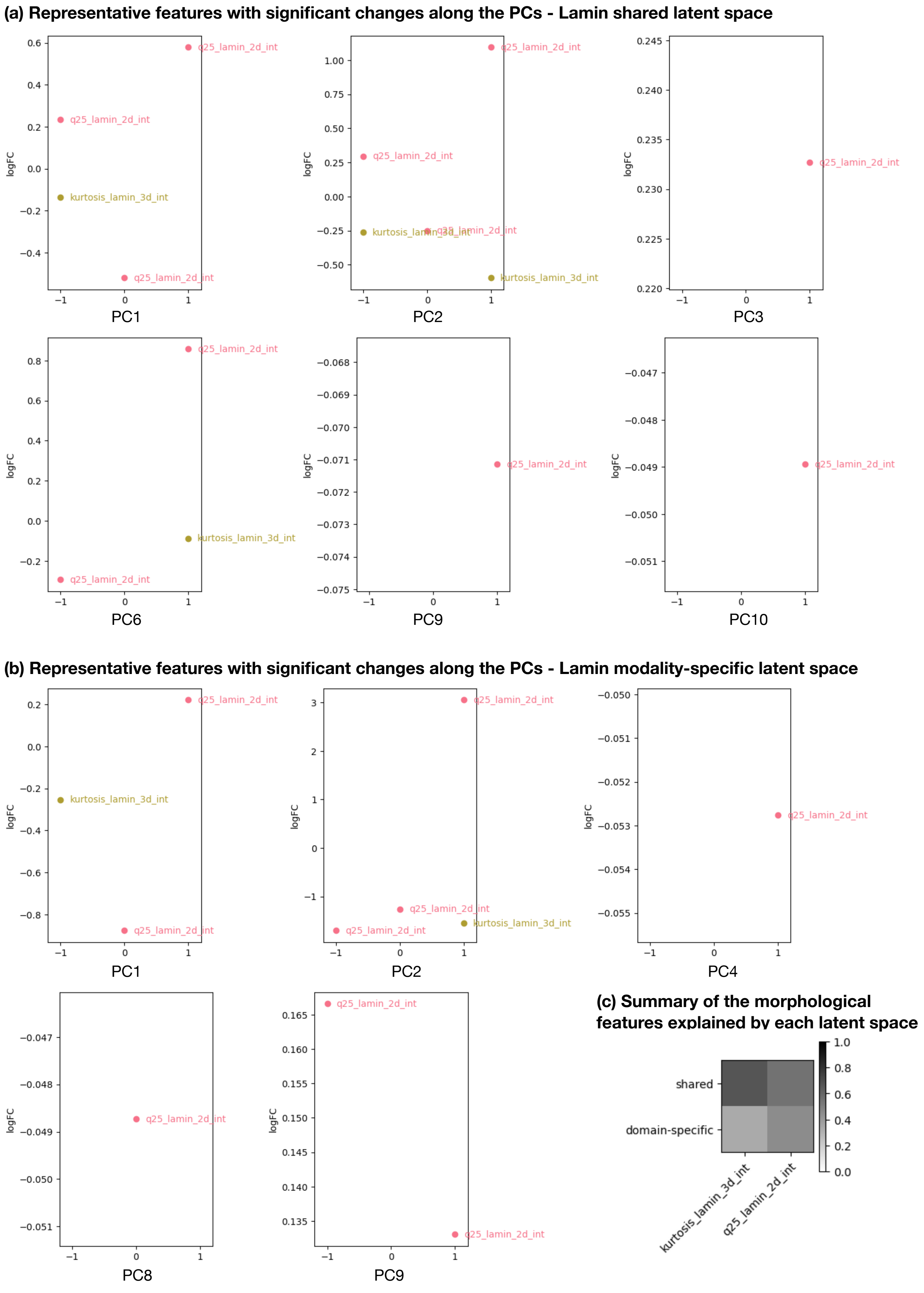

(a) Representative features with significant changes along the PCs - Lamin shared latent space
logFC
q25_lamin_2d_int
kurtosis_lamin_3d_int
PC1
PC2
PC3
PC6
PC9
PC10
(b) Representative features with significant changes along the PCs - Lamin modality-specific latent space
PC1
PC2
PC4
PC8
PC9
(c) Summary of the morphological features explained by each latent space
shared
domain-specific
kurtosis_lamin_3d_int
q25_lamin_2d_int


Figure S19

**Figure S19. Representative features in the lamin shared and modality-specific latent spaces.** Only one feature from each group of highly correlated features is plotted. The approach to feature grouping and selection of representative features is described in Extended Methods section. As described in the Extended Methods section, we consider three groups of cells for each $PC_i$ that are all at the center of other PCs, which are labeled as -1, 0, and 1 respectively.

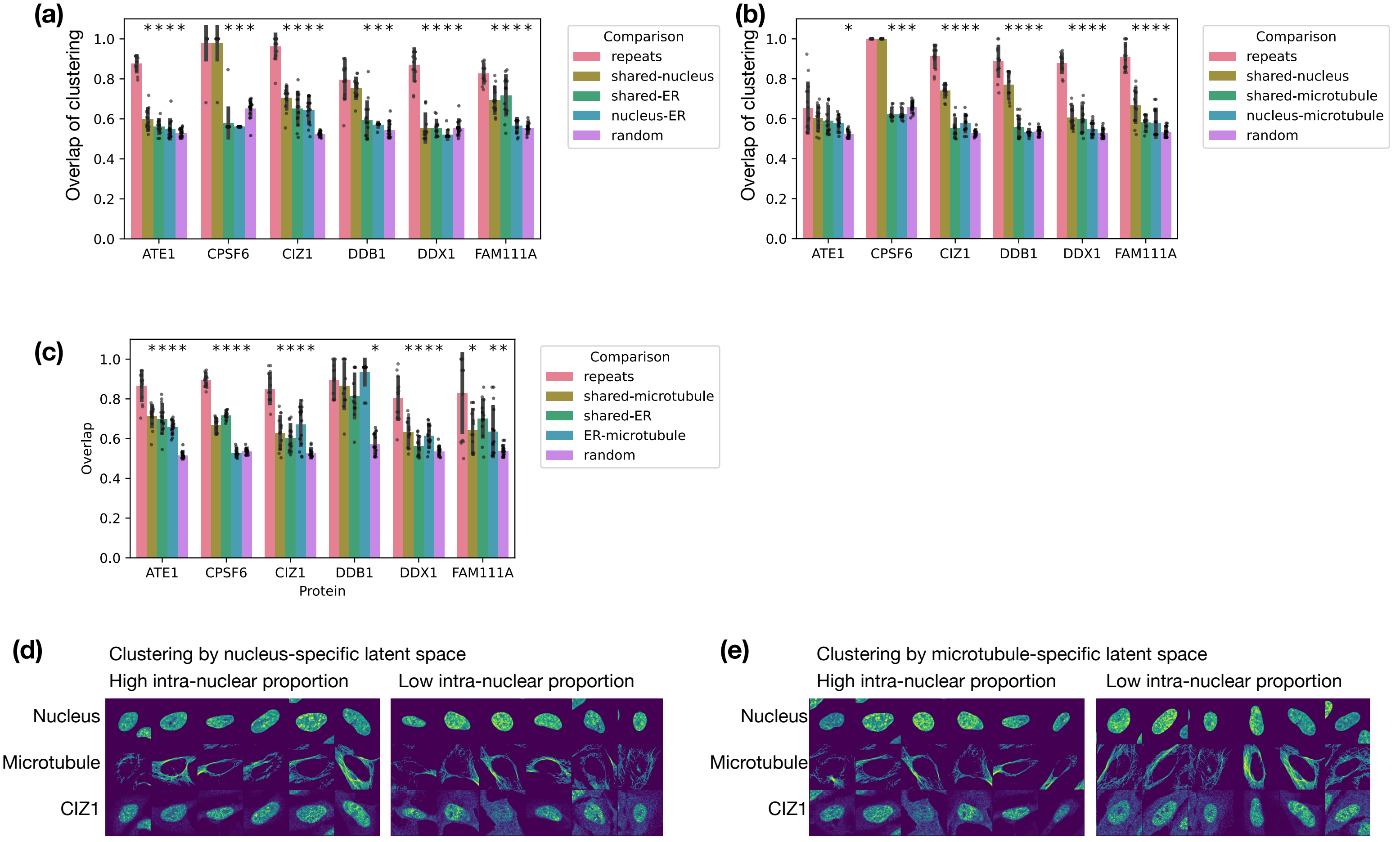
(a)
Overlap of clustering
1.0
0.8
0.6
0.4
0.2
0.0
ATE1
CPSF6
CIZ1
DDB1
DDX1
FAM111A
Comparison
repeats
shared-nucleus
shared-ER
nucleus-ER
random
(b)
Comparison
repeats
shared-nucleus
shared-microtubule
nucleus-microtubule
random
(c)
Overlap
Protein
Comparison
repeats
shared-microtubule
shared-ER
ER-microtubule
random
(d)
Clustering by nucleus-specific latent space
High intra-nuclear proportion
Low intra-nuclear proportion
Nucleus
Microtubule
CIZ1
(e)
Clustering by microtubule-specific latent space
High intra-nuclear proportion
Low intra-nuclear proportion
Nucleus
Microtubule
CIZ1

**Figure S20**

**Figure S20.**

(a)-(c) Fraction of overlapping cells between 1) repeated clustering of the shared latent space with different random initialization (repeats); 2) clustering of the shared and nucleus-specific latent spaces (shared-nucleus); 3) clustering of the shared and ER-specific latent spaces (shared-ER); 4) clustering of the nucleus-specific and ER-specific latent spaces (nucleus-ER); 5) random separation of cells into two groups (random). *: p-value $< 0.05$ compared to repeats. Panel (a) shows the model trained using nucleus and ER images. Panel (b) shows the model trained using nucleus and microtubule images. Panel (c) shows the model trained using microtubule and ER images.

(d) Examples of cells stained for CIZ1 in each cluster of the nucleus-specific latent space.

(e) Examples of cells stained for CIZ1 in each cluster of the microtubule-specific latent space.
